# RBMX enables productive RNA processing of ultra-long exons important for genome stability

**DOI:** 10.1101/2020.10.09.333039

**Authors:** Sara Luzzi, Gerald Hysenaj, Chileleko Siachisumo, Kathleen Cheung, Matthew Gazzara, Katherine James, Caroline Dalgliesh, Mahsa Kheirollahi Chadegani, Ingrid Ehrmann, Graham R Smith, Simon J Cockell, Jennifer Munkley, Yoseph Barash, David J Elliott

**Affiliations:** Biosciences Institute, Faculty of Medical Sciences, Newcastle University, Newcastle, United Kingdom; Bioinformatics Support Unit, Faculty of Medical Sciences, Newcastle University, Newcastle, United Kingdom; Department of Genetics, Perelman School of Medicine, University of Pennsylvania, Philadelphia, United States; Department of Biochemistry and Biophysics, Perelman School of Medicine, University of Pennsylvania, Philadelphia, United States; Department of Applied Sciences, Northumbria University, Newcastle, United Kingdom; Department of Computer and Information Science, University of Pennsylvania, Philadelphia, United States

## Abstract

Previously we showed that the germline-specific RNA binding protein RBMXL2 is essential for male meiosis where it represses cryptic splicing patterns (1). Here we find that its ubiquitously expressed paralog RBMX helps underpin human genome stability by preventing non-productive splicing. In particular, RBMX blocks selection of aberrant splice and polyadenylation sites within some ultra-long exons that would interfere with genes needed for normal replication fork activity. Target exons include within the *ETAA1* (*Ewings Tumour Associated 1*) gene, where RBMX collaborates with its interaction partner Tra2β to enable full-length exon inclusion by blocking selection of an aberrant 3’ splice site. Our data reveal a novel group of RNA processing targets potently repressed by RBMX, and help explain why RBMX is associated with gene expression networks in cancer, replication and sensitivity to genotoxic drugs.

## Introduction

Genome stability is essential to both prevent cancer and enable normal development (1). Nuclear RNA binding proteins can contribute to genome stability by regulating expression of genes involved in DNA replication and repair and/or directly participating in the DNA damage response (2). Furthermore, RNA binding proteins can suppress R-loop formation caused by aberrant excision of introns, which can lead to both transcription-replication conflicts and single-strand DNA damage (2).

Network analysis of alternative isoform ratios from thousands of tumours identified the nuclear RNA binding protein RBMX as a molecular switch closely linked to important cancer drivers (3). While the precise networks of gene expression controlled by RBMX in cancer cells are poorly understood, RBMX has also been identified as a potential tumour suppressor in oral and lung cancer, with tobacco induced mutations in *RBMX* predisposing smokers to future lung cancer development (4–8). RBMX also acquires somatic mutation in several cancer cohorts, including breast and endometrial cancer (9). Loss of the RBMX gene predisposes vemurafenib-resistant thyroid cancers to chromosome abnormalities (10). RBMX also contributes to mitotic progression and sister chromatid cohesion (11,12), and is required for normal brain development (13,14).

Data supports a direct role for RBMX in preventing DNA damage occurring at DNA replication forks stalled at repetitive DNA sequences (15,16). During replication stress, Replication Protein A (RPA) binds to single stranded DNA at stalled replication forks, leading to ATR activation by parallel pathways that depend on the protein kinases ETAA1 and TOPB1 respectively (17). RBMX binding to repetitive DNA sequences helps to stabilise TOBP1 to facilitate ATR activation at stalled replication forks, and depletion of RBMX causes replication defects and genome instability (15). RBMX protein also physically binds to the *NORAD* long ncRNA that is involved in DNA damage repair, although subcellular localisation experiments suggest that this direct protein-RNA association might not contribute to DNA repair (18,19). In addition, RBMX is needed for efficient p53-dependent DNA repair via non-homologous end joining (20), and depletion of RBMX sensitises U2OS osteosarcoma cells to DNA damage caused by ionising radiation and genotoxic drugs including cisplatin (21).

A critically important yet relatively unexplored molecular mechanism through which RBMX could promote genome stability is through splicing control. Splicing is a key process that allows maturation of protein-coding precursor RNAs (pre-mRNAs). In fact, most human genes are split up into exons and intervening intron sequences. Exons within pre-mRNAs are spliced together by the spliceosome to create mRNAs (22). Alternative splicing of exons in different orders allows production of several transcript and protein isoforms from the same genes (23,24). Although this promotes diversity and functional differentiation, deregulation of alternative splicing patterns are often associated with human pathologies including cancer (3,25). RBMX protein contains an N-terminal RNA Recognition Motif (RRM) and a C-terminal disordered region. A global search of exon skipping patterns in HEK293 controlled by RBMX showed it can directly promote splicing by recognising N6-methyladenosine (m6A) modification patterns (26,27). These patterns are deposited within pre-mRNAs by the N6 methyltransferase complex, which comprise a heterodimer of METTL3 and METTL14 methyltransferase-like proteins (27). RBMX also directly interacts with and frequently antagonises the splicing activity of the SR protein-family splicing regulator Tra2β (28–31). Tra2β normally operates as a splicing activator protein by promoting exon inclusion. The physiological importance of the antagonistic splicing complexes formed between Tra2β and RBMX are poorly understood. Interestingly, Adamson et al (21) speculated that RBMX may be important for splicing of genes involved in DNA repair, and demonstrated an RBMX-requirement for BRCA2 protein expression. BRCA2 is a key tumour suppressor that is involved in the homologous recombination pathway used to repair DNA damage, although the molecular connection between BRCA2 and RBMX needs to be identified.

Recent analyses from our group (32) show that a protein very similar to RBMX, called RBMXL2, reduces spliceosome selection of a group of weak splice sites, including previously unannotated “cryptic splice sites”. *RBMXL2* was derived by retrotransposition from the *RBMX* gene ~65 million years ago, and encodes a testis-specific protein with 73% sequence homology to RBMX (31,33). Genetic knockout of the *RBMXL2* gene in mice caused aberrant mRNA processing during meiosis, including the insertion of cryptic exons and premature terminal exons, and the modification of exon lengths through use of alternative splice sites. Furthermore, we showed that RBMXL2 is important for processing of some unusually large exons of over 1 kb in length. Intriguingly, one of the long exons controlled by RBMXL2 protein during meiosis was exon 11 of the mouse *Brca2* gene, where RBMXL2 repressed a cryptic 5’ splice site (32). The similarities between RBMX and RBMXL2 suggests the possibility that RBMX may play a similar role in mRNA processing in somatic cells which do not express RBMXL2, and this might possibly contribute to the phenotype of cells when RBMX is depleted (21).

Here we have used a global approach based on RNA sequencing to test this hypothesis in human breast cancer cells, and find that RBMX ensures correct mRNA processing and expression of genes that are key for genome stability. Importantly, our study provides molecular insights into how ultra-long exons are processed during RNA maturation.

## Results

### Global identification of a novel panel of RBMX-regulated RNA processing events

The sequence similarity between RBMX and RBMXL2 (34) prompted us to hypothesise that RBMX might control mRNA processing of genes involved in cell division and DNA damage response in somatic cells. To test this, we depleted *RBMX* from MDA-MB-231 cells (that model triple negative breast cancer) using siRNA, and performed RNA sequencing (RNA-seq) (Figure 1 – Figure supplements 1A, B). Western blot analysis confirmed >90% reduction of RBMX protein levels compared to control (Figure 1 – Figure supplement 1A), while RNA-seq analysis indicated a fold-change reduction in *RBMX* RNA levels upon treatment with RBMX siRNA of 0.12 compared to control, confirming successful RBMX knock-down.

In order to detect a wide range of transcriptome changes in RBMX targets we analysed our RNA-seq data using two bioinformatics programme, SUPPA2 and MAJIQ. SUPPA2 uses estimates of whole isoforms expression to detect global changes in RNA processing patterns (35). SUPPA2 analysis predicted 6708 differentially processed RNA isoforms upon RBMX knock-down. Strikingly, Gene Ontology (GO) analysis revealed that approximately 15% of the significantly enriched pathways were related to DNA replication, DNA repair and cell division, while others involved RNA processing, cellular response to stress and other stimuli (Figure 1A and Figure 1 – Source Data 1). MAJIQ is able to detect local splicing variations (LSV), including complex variations (LSV involving more than two alternative junctions), and de-novo variations (those involving unannotated junctions and exons) from RNA-seq data (36), thus providing complementary information to SUPPA2’s analysis.

**Figure 1.**
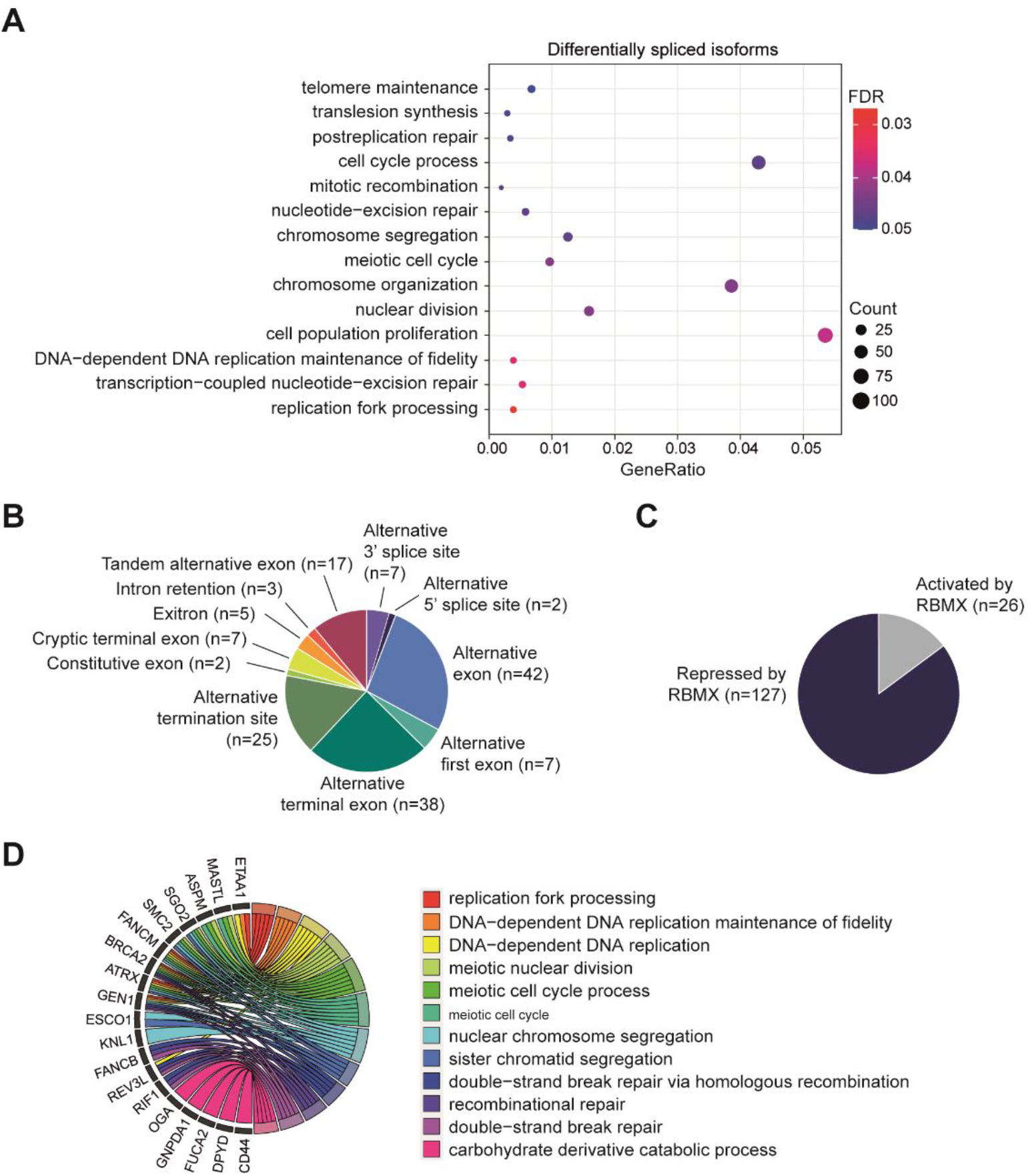
**(A)** Dot plot representing some of the most significantly enriched terms identified by Gene Ontology enrichment analysis of genes that undergo differential mRNA processing upon depletion of RBMX as detected by SUPPA2 analysis (35) (Figure 1 – Source Data 1). This plot was generated using GOstats v.2.54.0 (40) and ggplot2 (41) packages on R v.4.0.2. FDR, False discovery rate. Count, number of genes. **(B-C)** Pie charts representing types of the strongest mRNA processing defects detected after RBMX knock-down MDA-MB-231 cells by SUPPA2/MAJIQ analyses (35,36) (B), and whether these RBMX-controlled mRNA processing patterns are either repressed or activated by RBMX (C) (Figure 1 – Source Data 2). **(D)** Chord diagram presenting gene ontology of the most strongly RBMX-regulated targets identified by SUPPA2/MAJIQ (35,36) (Figure 1 – Source Data 2,3), produced using the Bioconductor GOplot (v1.0.2) package (42). Biological process GO terms with count > 4 and size < 250 (GO:0031297 and GO:0045005) are shown.

We also analysed RNA-seq data using the Majiq bioinformatic tool, which detects local splicing variations from RNA-seq data (36) and to identify the RNA processing patterns that most strongly depend on RBMX we visually inspected the RBMX predicted targets on the IGV genome browser (37). This visual search detected 155 strong changes in RNA processing including splice site selection, differential selection of terminal exons and alternative polyadenylation (polyA) sites, in addition to exon skipping (Figure 1B and Figure 1 – Source Data 2). Most of these RNA processing events (80%) were predicted to be repressed by RBMX (Figure 1C). Importantly, comparison with publicly available RNA-seq data (26) showed that while 48% of the same splicing events that we identified in MDA-MB-231 cells also switched mRNA processing after RBMX depletion in HEK293 cells (Figure 1 – Figure supplement 1C), these largely did not respond to depletion of either *METTL3* or *METTL14* m6A methyltransferases (Figure 1 – Figure supplement 1D). This indicates that the role of RMBX in repressing utilisation of splice sites is both cell-type and m6A-independent. Overall, 77% of the transcript variants that strongly changed after RBMX depletion were already annotated as mRNA isoforms on Ensembl (v94), and 23% were novel to this study (Figure 1 – Figure supplement 1E). Many genes contained single strong RBMX-regulated processing events, but in some genes like *CD44* and *TNC* (38,39) several adjacent exons are repressed by RBMX (Figure 1 – Source Data 2). Gene Ontology (GO) enrichment analysis of genes with strongly defective RNA processing patterns after RBMX depletion identified replication fork processing (GOBPID: 0031297, adjusted p-value = 3.94e-06), and DNA-dependent DNA replication maintenance of fidelity (GOBPID: 0045005, adjusted p-value = 1.32e-05) as the only significantly enriched terms (Figure 1D, Figure 1 – Figure supplement 1F and Figure 1 – Source Data 3). This indicates that RBMX controls RNA processing of genes involved in genome maintenance.

**Figure 1 – Figure Supplement 1.**
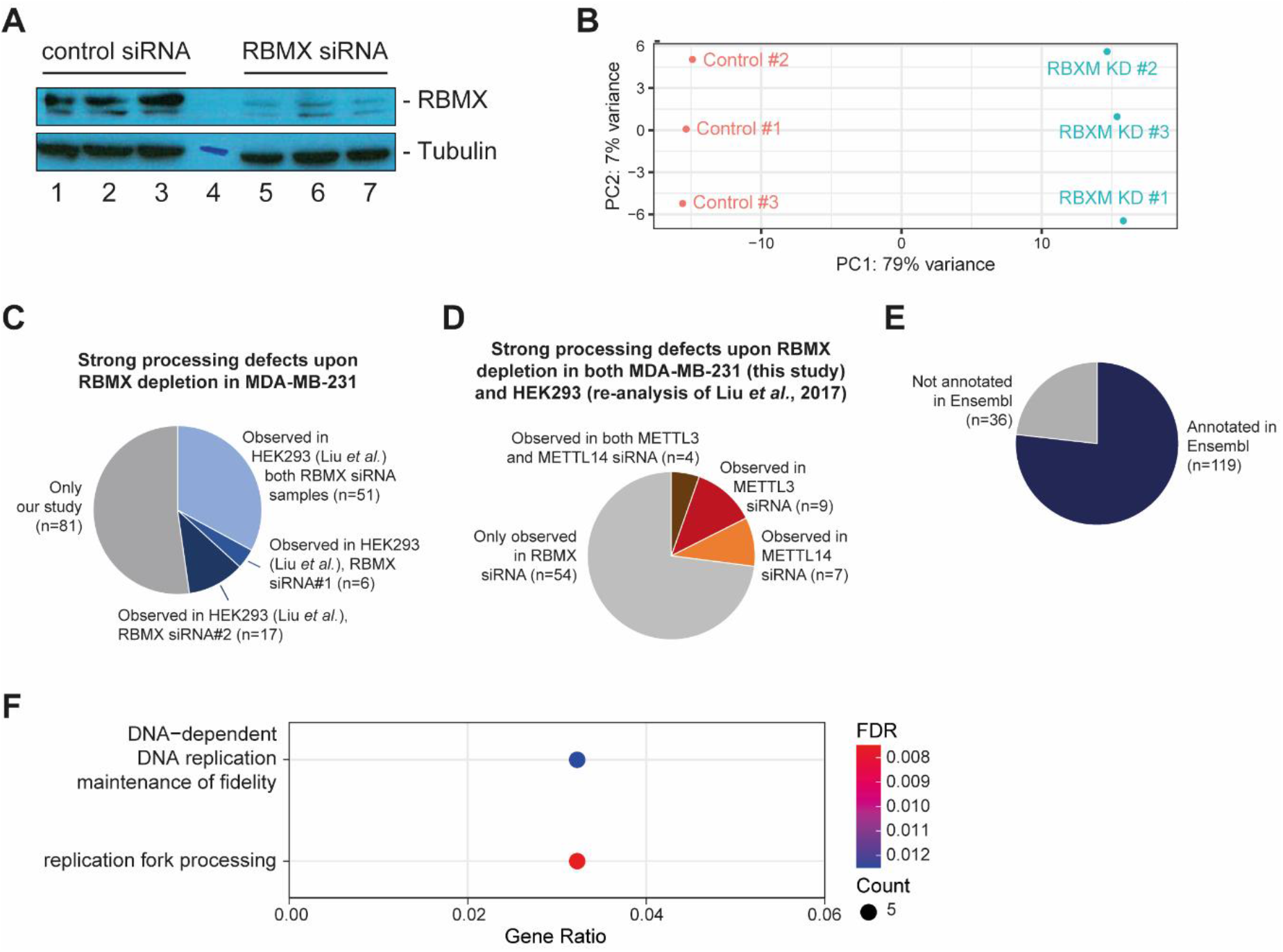
**(A)** Western blot analysis confirming reduction in RBMX protein levels in MDA-MB-231 cells after siRNA-mediated depletion of *RBMX*. Lanes 1-3, cells treated with control siRNA. Lane 4, molecular weight size marker. Lanes 5-7, cells treated with siRNA against RBMX. Samples were separated in the same gel and immunoblotted sequentially. Total RNA from the same cell samples was sequenced by RNA-seq**. (B)** Principal component analysis of RNA-seq data from cells treated with either control siRNA or siRNA against RBMX produced with DESeq2 v.1.16.1 (43) on R v.3.5.1. **(C)** Pie chart representing the proportion of the strongest RBMX-dependent RNA processing events that were identified in MDA-MB-231 cells (Figure 1 – Source Data 1) but could also be observed in RBMX-depleted HEK293 cells from (26), in either one or both samples treated with two separate siRNA against RBMX (RBMX siRNA#1 and #2). **(D)** Pie chart representing RBMX-dependent RNA processing events observed upon RBMX depletion in both MDA-M-231 and HEK293 cells (26) (see (C), blue slices) that can be detected after treatment with siRNA against either *METTL3* or *METTL14* or both. Data from (26). **(E)** Pie chart representing whether the strongest mRNA processing patterns detected after RBMX knock-down MDA-MB-231 cells by SUPPA2/MAJIQ analyses (35,36) (Figure 1 – Source Data 1) were already annotated in Ensembl (v94). **(F)** Dot plot representing gene ontology enrichment analysis of RBMX preferential targets identified by SUPPA2/MAJIQ (35,36) (Figure 1 – Source Data 2,3) generated using GOstats v.2.54.0 (40) and ggplot2 v.3.3.2 (41) packages on R v.4.0.2. FDR, False discovery rate; Count, number of genes.

### Splicing control by RBMX is required for normal expression of ETAA1 (Ewing’s Tumour Associated 1) protein kinase

The above data showed that genes involved in replication fork activity were globally enriched amongst genes showing strong splicing changes after RBMX depletion. Replication fork accuracy is critical for genome stability, and one of the most strongly RBMX-dependent RNA processing patterns was for the *ETAA1 (Ewing’s Tumour-Associated Antigen 1*) gene. *ETAA1* encodes a protein essential for replication fork integrity and processivity (44–46) (Figure 1D). Depletion of RBMX protein in MDA-MB-231 cells dramatically changed the *ETAA1* splicing profile, increasing selection of a very weak 3’ splice site within exon 5 (Weight Matrix Model score: −1.67, compared to 9.11 for the stronger upstream 3’ splice site) (Figure 2A and Figure 2 – Figure Supplement 1A). We experimentally confirmed this *ETAA1* splicing switch using duplex RT-PCR from cells depleted for RBMX. While control cells had almost total inclusion of the full-length version of *ETAA1* exon 5, there was a significantly increased selection of the exon 5-internal 3’ splice site in the RBMX depleted cells (p=0.099, Figures 2B, C). We also found that RBMX-mediated splicing control of *ETAA1* exon 5 is not specific to MDA-MB-231 cells. Analysis of an RNA-seq dataset made from HEK293 cells from which RBMX had been depleted with two independent siRNAs (26) demonstrated a similar switch in the *ETAA1* splicing profile, while also detecting RBMX-mediated repression of an additional weak 3’ splice site that is infrequently used in MDA-MB-231 cells (Figure 2 – Figure Supplement 1B).

**Figure 2.**
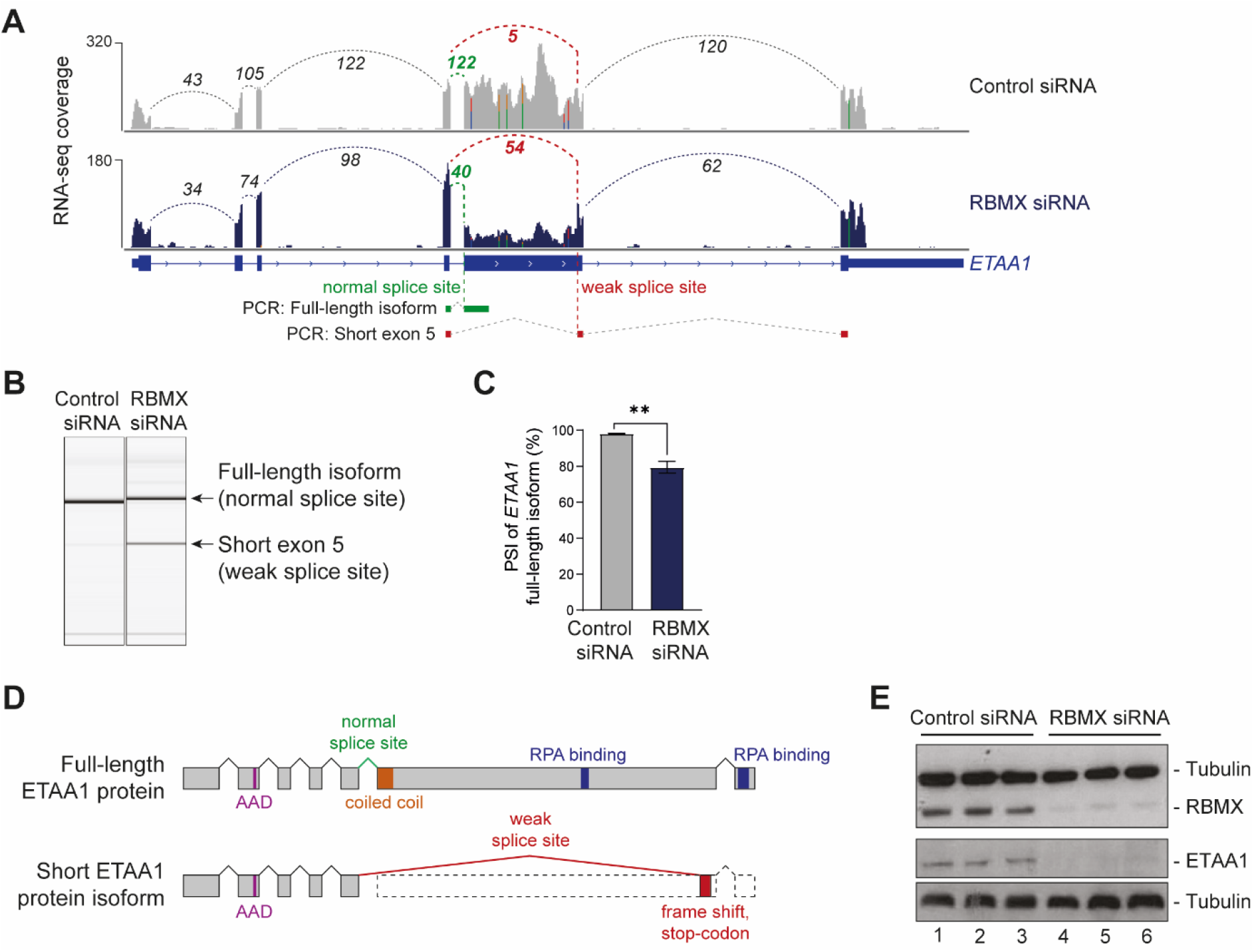
RBMX is essential for normal ETAA1 protein expression. **(A)** Snapshot from IGV browser (37) showing merged RNA-seq tracks over the *ETAA1* gene, from triplicate MDA-MB-231 cells taken after either *RBMX* depletion (“RBMX siRNA”) or control treatment (“Control siRNA”). Splice junctions between *ETAA1* exons are shown with dotted lines. The splice junction mainly used in controls cells is shown in green, and the splice junction normally used in RBMX-depleted cells is shown in red. The position of the multiplex PCR products of the analysis in **(B)** are also shown. Numbers indicate read count over each splice junction. **(B)** Representative capillary gel electrophoretograms showing RT-PCR validation of the changing *ETAA1* splicing pattern after siRNA-mediated depletion of *RBMX* in MDA-MB-231 cells. **(C)** Bar chart associated with the experiment shown in (B) shows Percentage Splicing Inclusion (PSI) of the full-length *ETAA1* isoform using data from three biological replicates. Bars, standard error. **, p-value < 0.01 as calculated by t-test with Welch’s correction. **(D)** Schematic representation of the ETAA1 protein resulting from either normal splicing (Full-length) or aberrant splicing detected upon RBMX depletion (Short isoform). AAD, ATR-activation domain; RPA, Replication Protein A1 binding motif. **(E)** Western blot analysis confirming no full-length ETAA1 protein is detected after RBMX depletion. The Western blot contains 3 replicate protein samples for RBMX depletion (“RBMX siRNA”) or after treatment with a control siRNA (“Control siRNA”). The same samples ran on two different gels and were probed sequentially probed with either anti-RBMX and anti-tubulin antibodies, or with anti-ETAA1 and anti-tubulin antibodies.

The full-length ETAA1 protein is 926 amino acids long. Splicing selection of the *ETAA1* exon 5-internal 3’ splice site produces an mRNA isoform predicted to encode an ETAA1 protein isoform of just 202 amino acids (Figure 2D). Although the *ETAA1* exon 5-internal 3’ splice site is annotated on Ensembl (v94), it is rarely selected in cells treated with control siRNAs (Figure 2A). Confirming that correct expression of ETAA1 protein depends on RBMX, Western blot analysis with an antibody specific to ETAA1 protein showed strong reduction of the full-length ETAA1 protein after RBMX depletion (Figure 2E). Such a short ETAA1 protein would lack RPA binding motifs (Figure 2D) and thus be unable to operate similarly to the full-length ETAA1 protein isoform. Hence normal *ETAA1* gene function relies on the RNA processing activity of RBMX.

**Figure 2 – Figure Supplement 1.**
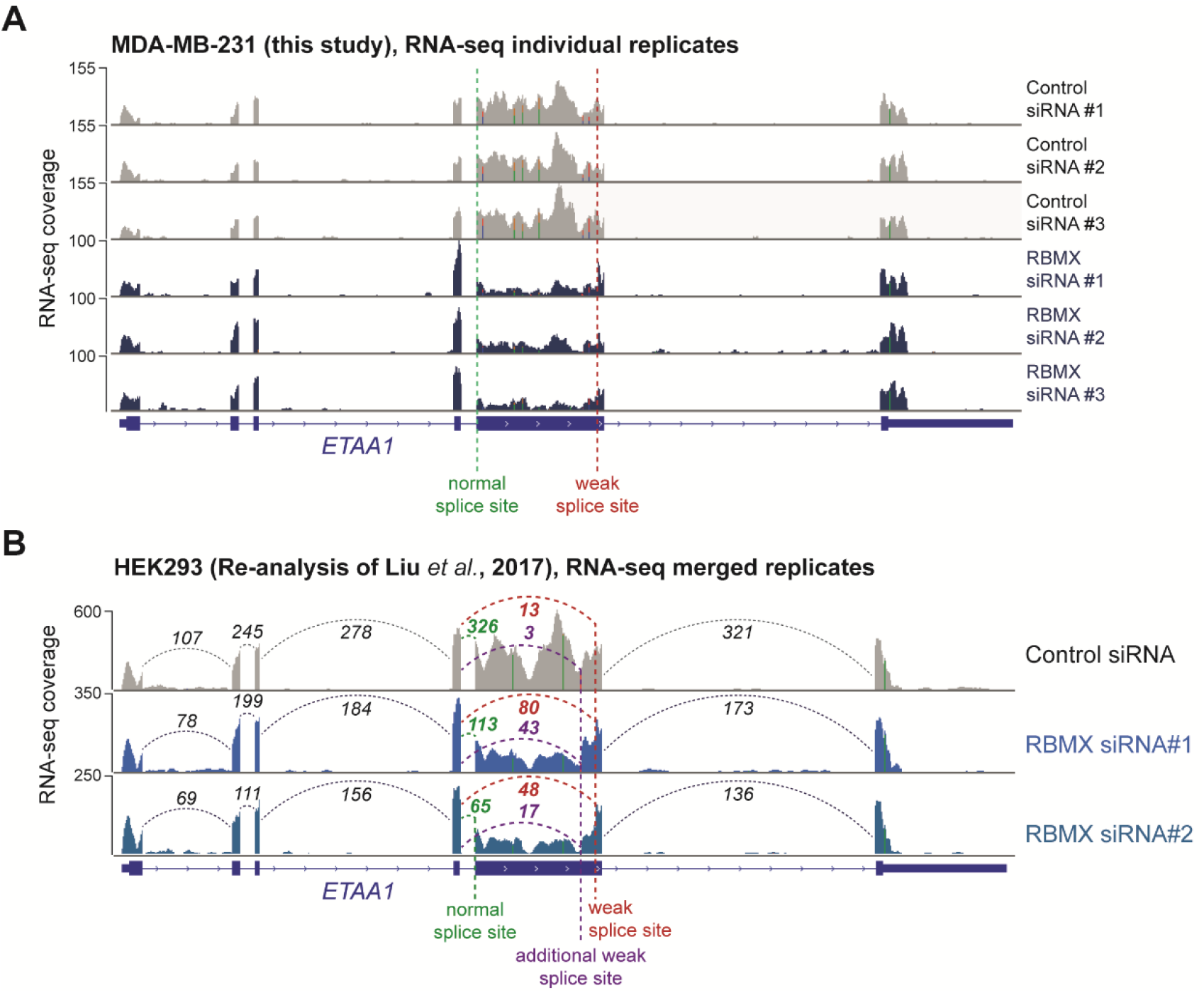
**(A)** Snapshot from the IGV browser (37) showing single replicate RNA-seq tracks from MDA-MB-231 cells treated with either control siRNA or siRNA against RBMX across the *ETAA1* gene. The two 3’ splice sites on *ETAA1* exon 5 are shown. **(B)** Snapshot from the IGV browser (37) over the *ETAA1* gene, showing merged RNA-seq tracks from HEK293 cells treated with either control siRNA (“Control siRNA”) or two separate siRNAs against *RBMX* (“RBMX siRNA1” and “RBMX siRNA2”). The strong upstream 3’ splice site and the two weak downstream 3’ splice sites on *ETAA1* exon 5 are shown. The HEK293 RNA-seq data is from (26).

### RBMX cooperates with Tra2β to suppress cryptic splicing within ETAA1 exon 5

Two possible mechanistic models could explain the different use of 3’ splice sites within *ETAA1* exon 5 in RBMX-depleted cells: RBMX could normally promote recognition of the strong upstream splice site; or, RBMX could normally prevent usage of the weak downstream splice site. In order to distinguish between these two possibilities, we performed a minigene assay. Briefly, a fragment of *ETAA1* exon 5 that spanned the weak internal 3’ splice site and flanking genomic regions (but not the stronger upstream 3’ splice site) was cloned into an expression plasmid between two β-globin exons (47) (Figure 3A and Figure 3 – Figure Supplement 1A). When transfected into HEK293 cells, this minigene expressed a splice variant including the shorter version of *ETAA1* exon 5 (Figure 3B). Co-transfection of RBMX only weakly suppressed inclusion of the short exon, evidenced by a slight but not significant production of an RNA isoform in which the β-globin exons are directly spliced together. (Figures 3B, C).

**Figure 3.**
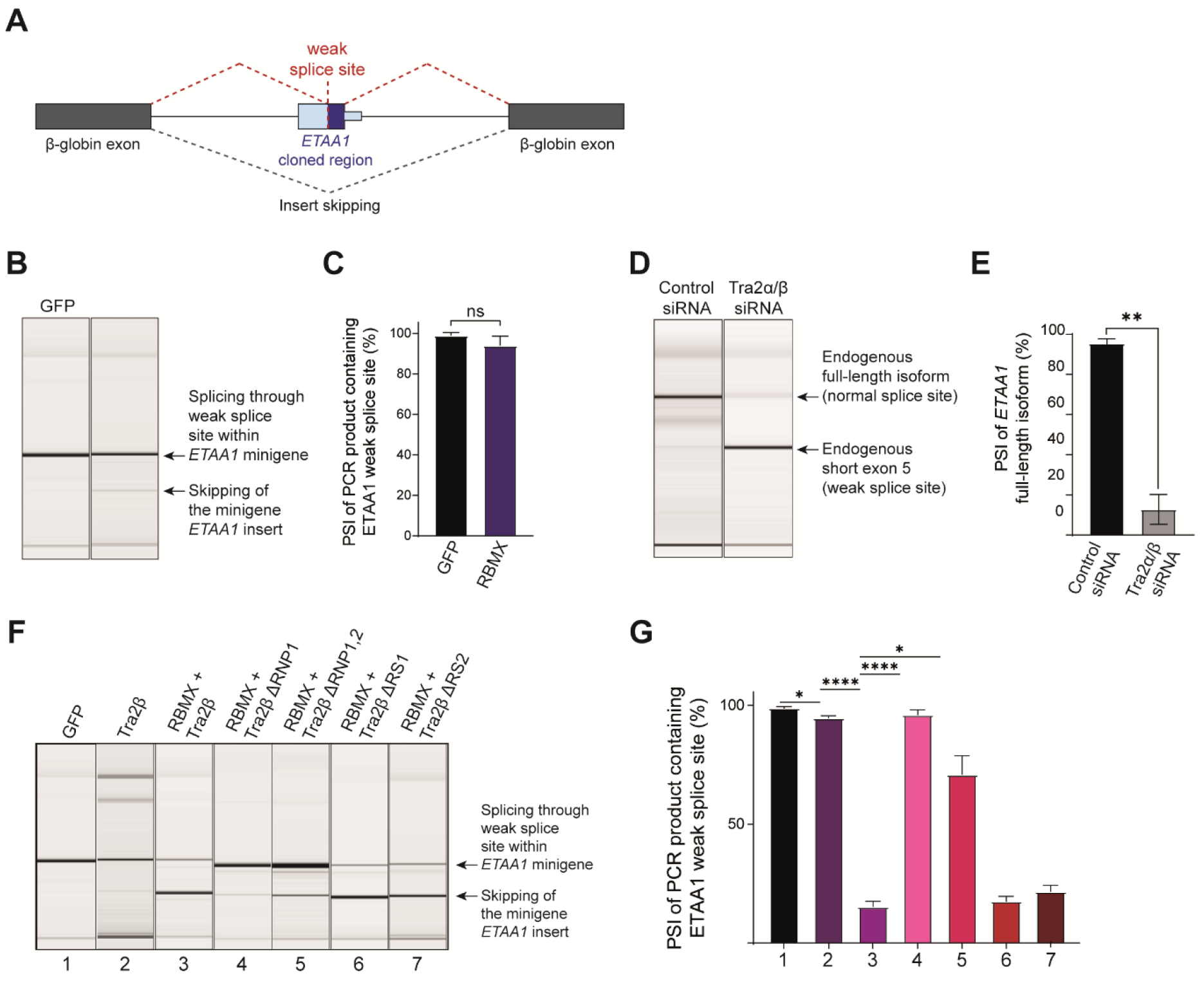
RBMX cooperates with Tra2β to enable splicing inclusion of full-length *ETAA1* exon 5. **(A)** Schematic representation of the minigene containing a portion of the *ETAA1* gene around the weak splice site used upon depletion of RBMX (see Figure 2A and Figure 3 – Figure Supplement 1A) cloned between the two β-globin exons of the pXJ41 plasmid (47). Dotted lines indicate the two possible splicing patterns of the *ETAA1* minigene. **(B)** Representative capillary gel electrophoretograms showing RT-PCR analysis of the *ETAA1* minigene transfected in MDA-MB-231 cells co-transfected with expression plasmids encoding either GFP or RBMX. The two products derived from either recognition of the *ETAA1* weak splice site (upper band) or skipping of the *ETAA1* insert (lower band) are indicated. **(C)** Bar chart associated with (B) shows Percentage of Splicing Inclusion (PSI) of the short form of the *ETAA1* exon (shown as dotted red lines in (A)). Bars, standard error. *, p-value < 0.05 as calculated by t-test with Welch’s correction across 4 biological replicates. **(D)** Representative capillary gel electrophoretogram showing RT-PCR validation of the changing *ETAA1* splicing pattern after joint depletion of Tra2α and Tra2β in MDA-MB-231 cells. **(E)** Bar chart associated with panel (D) showing Percentage Splicing Inclusion (PSI) of the full-length isoform of *ETAA1* and including data from three biological replicates. Bars, standard error. ****, p-value < 0.0001 as calculated by t-test with Welch’s correction. **(F)** Representative capillary gel electrophoretograms showing RT-PCR analysis of the *ETAA1* minigene co-transfected into MDA-MB-231 cells with expression plasmids encoding the indicated proteins and protein isoforms. Tra2βΔRNPI and Tra2βΔRNP1/2 are protein isoforms of Tra2β lacking either one or both RNP motifs within the RRM (50). Tra2βΔRSI and Tra2βΔRS2 are protein isoforms of Tra2β lacking one of its arginine-serine RS domains. The two products derived from either recognition of the *ETAA1* weak splice site (upper band) or skipping of the *ETAA1* insert (lower band) are indicated. **(G)** Bar chart associated with (F) shows Percentage of Splicing Inclusion (PSI) of the *ETAA1* spliced insert as shown in (A) (dotted red lines), including data from 3 to 7 biological replicates. Bars, standard error. *, p-value < 0.05 and ****, p-value < 0.0001 as calculated by t-test with Welch’s correction.

Co-transfection of an expression plasmid encoding RBMX only slightly repressed selection of the weak *ETAA1* splice site contained in the minigene. We thus further examined the mechanism of RBMX-dependent processing of *ETAA1* exon 5. RBMX directly interacts with the splicing regulator Tra2β, frequently to antagonise splicing activation (28–31). Previously published RNA-seq data indicated MDA-MB-231 cells that were jointly depleted for both Tra2β and its partially redundant paralogue Tra2α (48) had similar *ETAA1* exon 5 splicing defects to those observed upon *RBMX* depletion (Figure 3 – Figure Supplement 1B). Furthermore, Tra2β-RNA association analysis using iCLIP revealed that *ETAA1* exon 5 is also directly bound by Tra2β (48) (Figure 3 – Figure Supplement 1B). Consistent with a role for Tra2 proteins in regulating processing of *ETAA1* exon 5, RT-PCR analysis confirmed a switch in splicing inclusion from the long to the short version of *ETAA1* exon 5 in response to joint depletion of Tra2α and Tra2β in MDA-MB-231 cells (Figures 3D, E). We also obtained very similar results in the MCF7 breast cancer cell line (Figure 3 – Figure Supplements 1C, D). Thus Tra2-mediated repression of the internal weak 3’ splice site within *ETAA1* exon 5 occurs in multiple breast cancer cell types.

Further minigene experiments supported Tra2-mediated repression of *ETAA1* exon 5 internal splicing, and moreover an interaction between Tra2β and RBMX. Co-transfection into HEK293 cells of an expression vector encoding Tra2β with the *ETAA1* exon 5 minigene (Figure 3A) confirmed that splicing inclusion of the shorter *ETAA1* exon 5 was weakly but significantly repressed by Tra2β (Figures 3F, G). Strikingly, while Tra2β normally operates as a splicing activator (49), co-transfection of RBMX and Tra2β resulted in much stronger repression of the shorter version of *ETAA1* exon 5 than either RBMX or Tra2β alone (Figures 3F, G). Interestingly, co-transfection of a Tra2β isoform containing a deletion within its RRM domain (either Tra2βΔRNPI, or Tra2βΔRNPI,2) (50) was sufficient to efficiently reduce repression of the weak *ETAA1* splice site (Figures 3F, G). This suggests that direct Tra2β-RNA interactions are important for RBMX-mediated suppression of the short splice isoform of *ETAA1*. However, deletion of RS1 or RS2 domains (normally used for splicing activation by Tra2β) did not block splicing repression of the short ETAA1 exon.

**Figure 3 – Figure Supplement 1.**
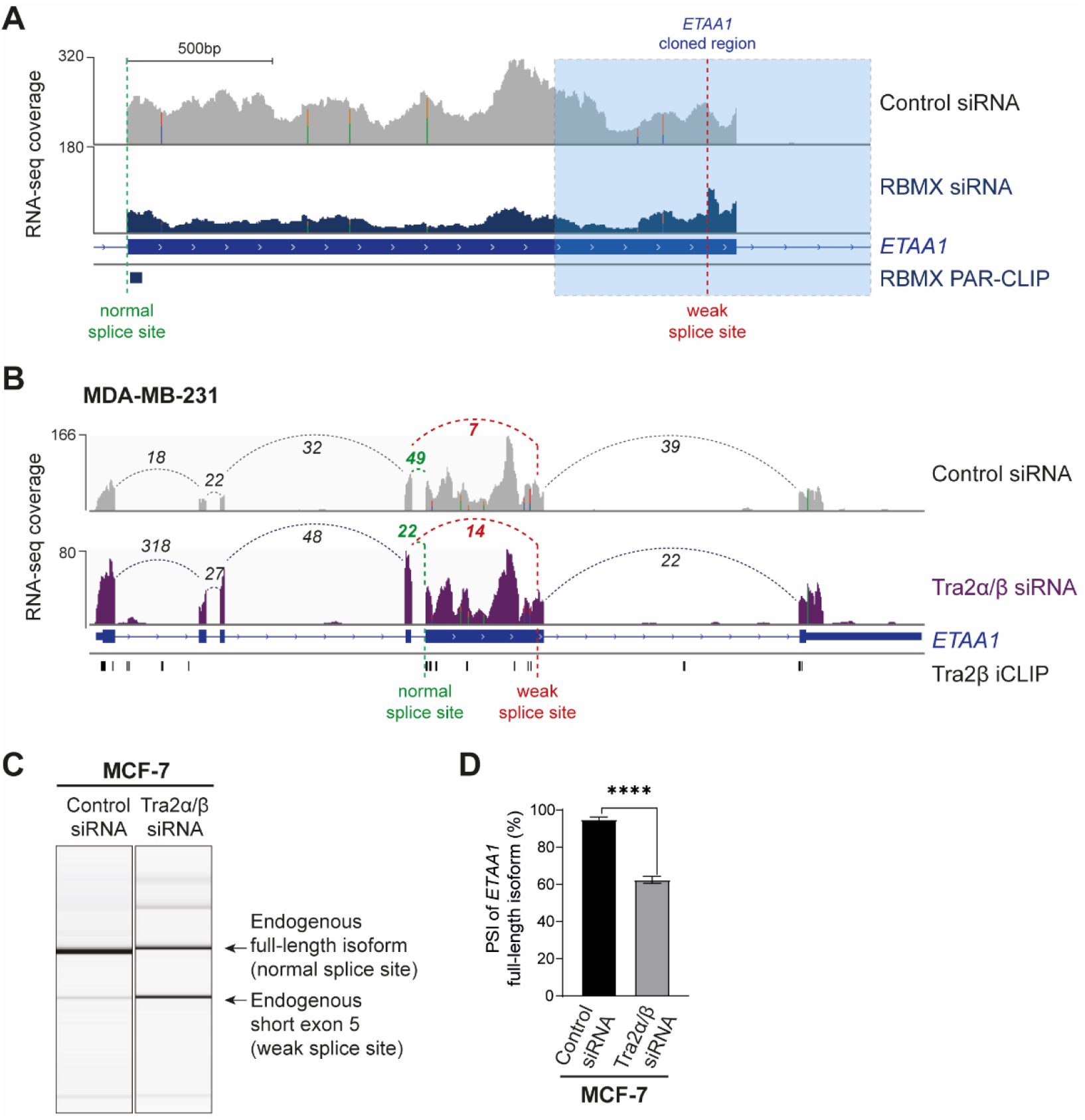
**(A)** Snapshot over the *ETAA1* exon 5 showing merged RNA-seq tracks from MDA-MB-231 treated with either control siRNA or siRNA against RBMX, as well as the RBMX PAR-CLIP binding site from (26). The two 3’ splice sites are shown. The region cloned into the pXJ41 minigene (47) (Figure 3A) is indicated in blue. **(B)** Snapshot over the full-length *ETAA1* gene shows merged RNA-seq tracks from MDA-MB-231 treated with either control siRNA (“Control siRNA”) or two siRNAs against *TRA2A* and *TRA2B* (“Tra2α/β siRNA”), and from iCLIP for Tra2β RNA binding. The two 3’ splice sites on *ETAA1* exon 5 are shown. RNA-seq and iCLIP data are from (48). **(C)** Representative capillary gel electrophoretograms showing RT-PCR validation of the changing *ETAA1* splicing pattern after joint depletion of Tra2α and Tra2β in MCF7 breast cancer cells and NCI-H520 lung cancer cells. **(D)** Bar chart associated with (C) showing Percentage Splicing Inclusion (PSI) of the full-length isoform of *ETAA1* and including data from three biological replicates. Bars, standard error. ****, p-value < 0.0001 as calculated by t-test. *, p-value < 0.5 as calculated by t-test.

### RBMX efficiently represses a spectrum of alternative RNA splice sites in ultra-long exons within genes that are important for genome stability

The above data showed that RBMX prevents cryptic mRNA processing of the ultra-long *ETAA1* exon 5, which at 2111nt is considerably longer than the 129 nt median size of human exons. Further examination revealed that RBMX also controls productive splicing patterns of ultra-long exons in other genes important in genome stability. RBMX efficiently represses a cryptic (defined as not annotated in Ensembl v94) 3’ splice site within the 4161nt exon 13 of the *REV3L* (*REV3 Like, DNA Directed Polymerase Zeta Catalytic Subunit*) gene (Figure 4A). *REV3L* encodes the catalytic component of DNA polymerase ζ that helps repair of stalled replication forks (51) and trans-lesion DNA replication. Furthermore, REV3L plays a key role in the response to ionising radiation, which is known to be defective in RBMX-depleted cells (21). The high amplitude splicing switch that occurs in response to RBMX depletion removes coding information for 1387 amino acids from the REV3L protein (normally 3130 amino acids long). Comparative analysis between our RNA-seq and published RNA-seq data from HEK293 cells (26) revealed the same RBMX-dependent and m6A-independent processing of *REV3L* pre-mRNA (Figure 4 – Figure Supplement 1A). Additionally, analysis of RBMX-RNA association by PAR-CLIP from HEK293 cells (26) revealed multiple direct RBMX binding sites to *REV3L* exon 13 (Figure 4A and Figure 4 – Figure Supplement 1A) supporting a direct role for RBMX in *REV3L* mRNA processing. RBMX also prevents the use of internal 3’ splice sites in ultra-long internal exons in the *RIF1* (*Replication Timing Regulatory Factor 1*) gene (3236nt exon 30, Figure 4 – Figure Supplement 1B) and *ASPM* (*Assembly Factor For Spindle Microtubules*) gene (4754nt exon 18, Figure 4 – Figure Supplement 1C).

**Figure 4.**
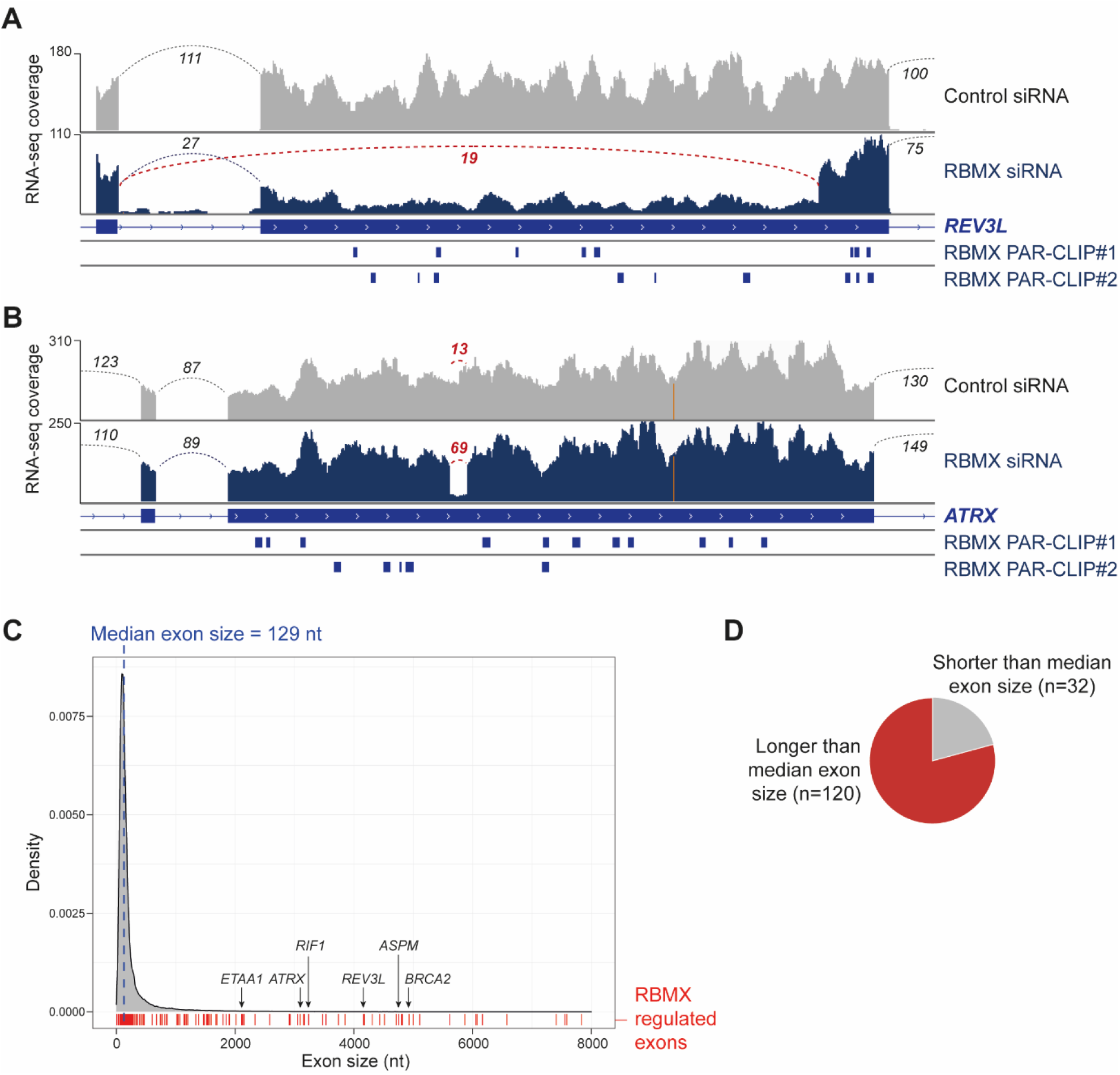
RBMX efficiently represses a spectrum of RNA splicing events within large exons in MDA-MB-231 cells. **(A-B)** Snapshots from IGV browser (37) over the *REV3L* and *ATRX* genes showing merged RNA-seq tracks from triplicate MDA-MB-231 cells taken after either *RBMX* depletion (“RBMX siRNA”) or control treatment (“Control siRNA”), as well as RBMX PAR-CLIP from (26). **(C)** Distribution of human exon sizes from genome build hg38 annotated in Ensembl (v101). Dashed blue line denotes the median exon size (129 nt). Exons that contain sites strongly regulated by RBMX (Figure 1 – Source Data 2) are indicated in red. Plot was created using ggplot2 (41) on R v.4.0.2. **(D)** Pie chart representing the proportion of exons containing RBMX-regulated RNA processing sites that are either larger (red slice) or shorter (grey slice) than the median size of human exons.

The RBMX-mediated repression of internal 3’ splice sites in the *ETAA1, REV3L* and *RIF1* genes enables production of full-length protein coding mRNAs that are important for genome stability (52). RBMX also controls splicing of an 87nt exitron (*i.e*. exonic intron) within the 3097nt exon 9 of the *ATRX* gene, which encodes a SWI/SNF family ATP-dependent chromatin remodeller involved in repair of stalled replication forks, gene regulation and chromosome segregation (53) (Figure 4B). Although annotated on Ensembl v.94 as an intron retention event, analysis of exon-junction reads show that this *ATRX* exitron is hardly ever used in control MDA-MB-231 cells (Figure 4B). Furthermore, PAR-CLIP data (26) confirmed that RBMX protein directly interacts with *ATRX* exon 9 at multiple sites (Figure 4B).

Strikingly, we found that most of the mRNA processing events that we identified to be strongly regulated by RBMX are located within large exons, 80% of which are longer than the 129 nt median exon size (Figures 4C, D and Figure 1 – Source Data 2). Consistently, analysis of gene expression patterns from HEK293 cells (26) showed that 60% of the exons with reduced splicing after RBMX depletion are also longer than the median human exon size (Figure 4 – Figure Supplements 1D, E). This is consistent with RBMX ensuring correct inclusion of unusually long exons during mRNA processing across different cell types.

**Figure 4 – Figure Supplement 1.**
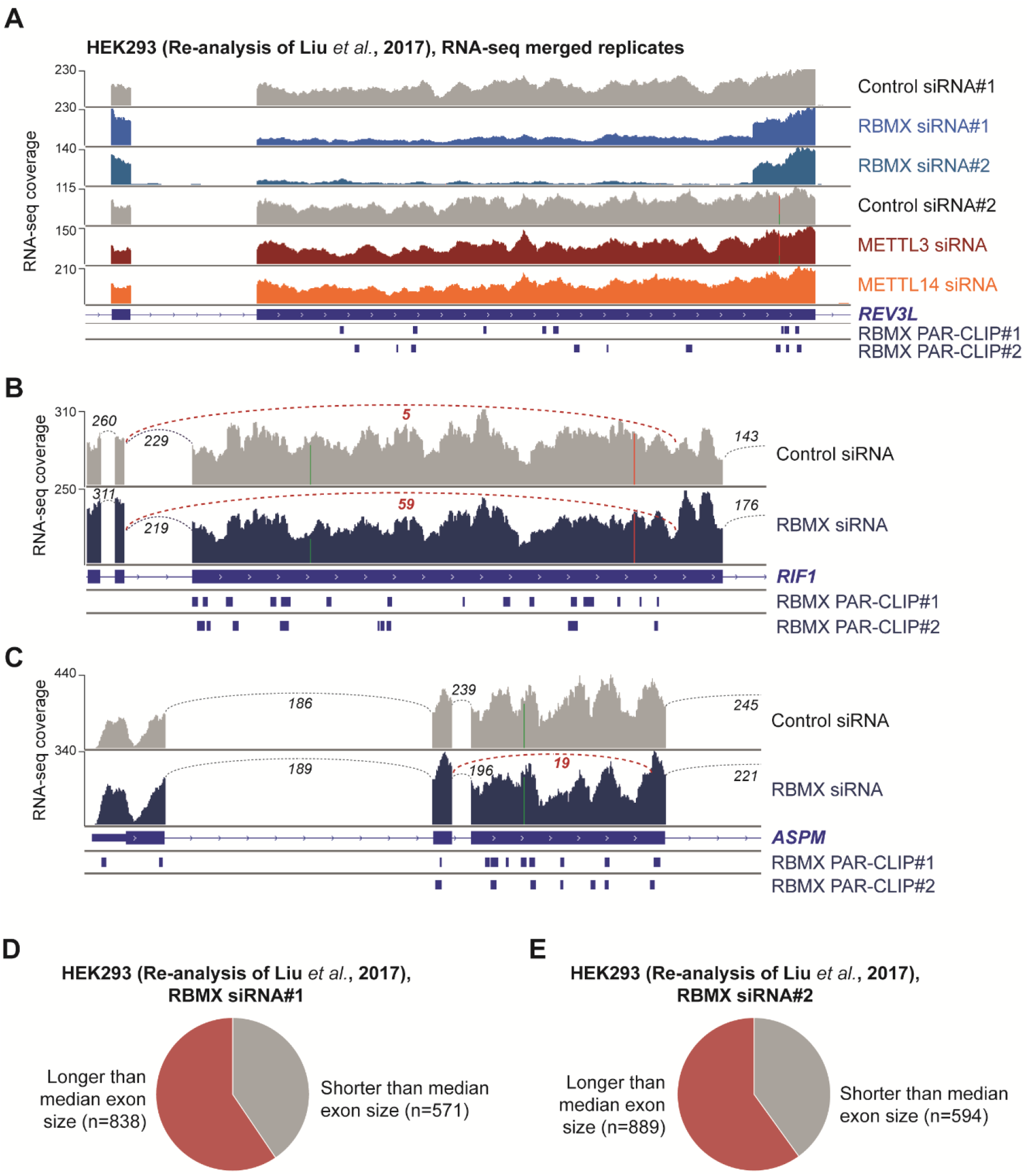
**(A)** Snapshot from the IGV browser (37) over *REV3L* exon 13, showing merged RNA-seq tracks from HEK293 cells treated with control siRNA (“Control siRNA#1” and “Control siRNA#2”), two separate siRNAs against *RBMX* (“RBMX siRNA#1” and “RBMX siRNA#2”), siRNA against METTL3 (“METTL3 siRNA”) and against METTL14 (“METTL14 siRNA”), as well as RBMX PAR-CLIP. RNA-seq data from (26). **(B-C)** Snapshots from IGV browser (37) over the *RIF1* and *ASPM* genes showing merged RNA-seq tracks from triplicate MDA-MB-231 cells taken after either *RBMX* depletion (“RBMX siRNA”) or control treatment (“Control siRNA”), as well as RBMX PAR-CLIP from (26). Splicing junctions between exons are represented with dotted lines. **(D-E)** Pie charts representing the proportion of exons that are downregulated upon RBMX depletion (DEXseq2 control/knock-down fold-change > 1) with two separate siRNAs in HEK293 cells and are either larger (red slice) or shorter (grey slice) than the median size of human exons. Data from (26).

### RBMX represses upstream transcriptional termination sites within key genes important for genome stability

Previous data have implicated RBMX with a role in splicing regulation (27). However, the above analyses additionally predicted that RBMX controls selection of transcription termination events for a panel of 64 genes (Figure 1 – Source Data 2 and Figure 5 – Source Data 1). Interestingly, *BRCA2* was identified as one of these differentially terminated mRNA transcripts (Figure 1D). Consistent with this, more detailed observation showed that RNA-seq reads asymmetrically mapped up to the first part of the ~5Kb long exon 11 of the *BRCA2* gene, with reduced read density for the remainder of this exon as well as all downstream exons (Figure 5A). Most of the premature termination events identified in our analysis occurred at termination sites previously mapped by high-throughput studies (54), while 7 events appeared to involve novel termination sites including within *BRCA2* (Figure 5B). Despite the lack of previously detected polyadenylation events at the site of the *BRCA2* exon 11 where RNA-seq reads dropped in RBMX-depleted cells, this genomic region does contain a canonical consensus sequence for polyadenylation (Figure 5C) (55–57). Furthermore, visual inspection of an alignment file that compares the RNA-seq reads from cells treated with RBMX siRNA to control siRNA using the bamcompare tool from deepTools v3.5.0 (58) confirmed reduction of RNA-seq reads after this putative polyA site upon RBMX depletion (Figure 5C). Consistent with this, RT-PCR analysis showed that the relative abundance of a PCR product spanning the premature termination site, normalised over a region upstream, was significantly reduced in RBMX-depleted cells compared to control (Figures 5D, E). This suggests that RBMX prevents premature transcription termination within *BRCA2* exon 11.

**Figure 5.**
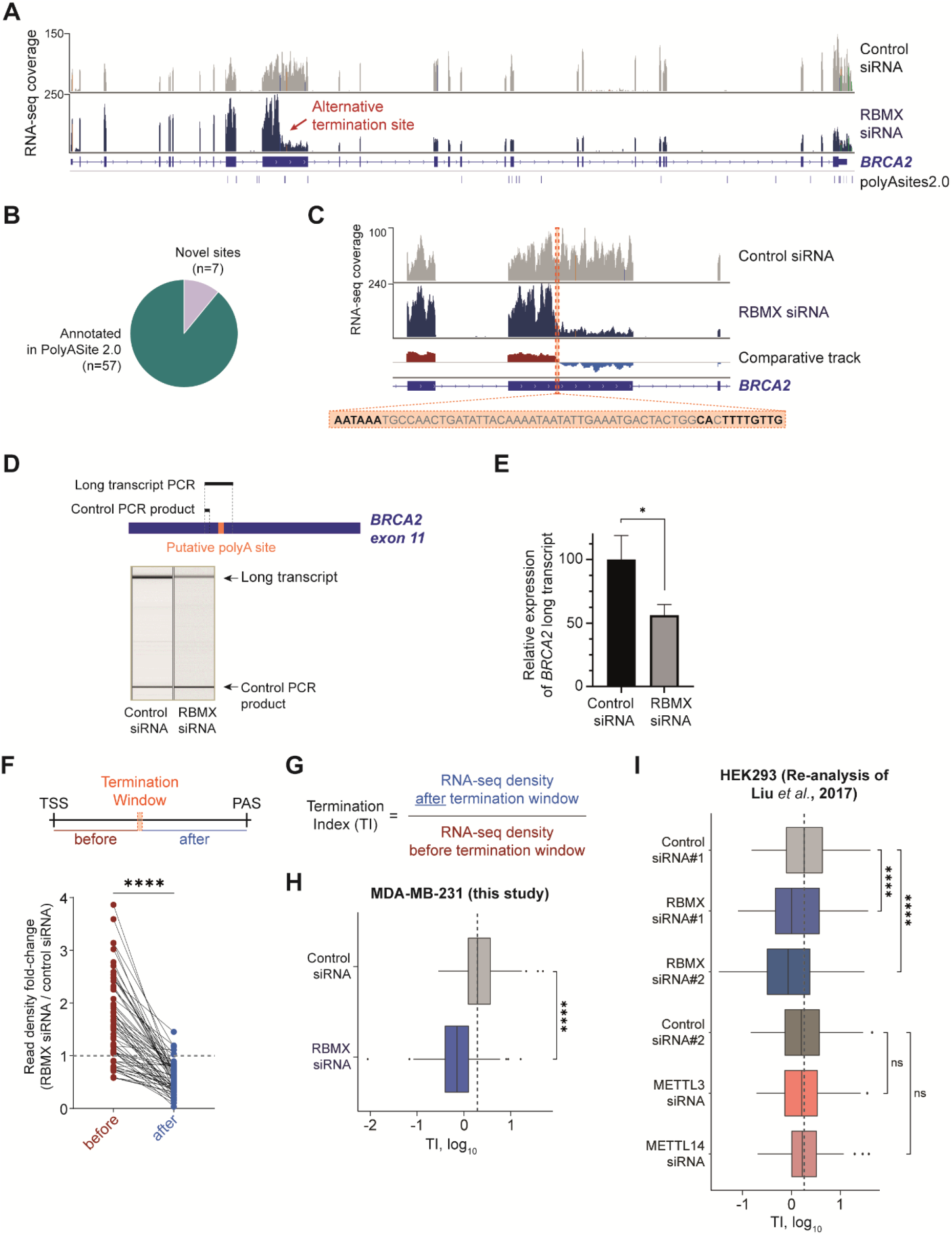
Depletion of RBMX leads to significantly strong reduction of full-length mRNA transcripts from genes involved in genome maintenance. **(A)** Snapshots from IGV browser (37) over the *BRCA2* gene showing merged RNA-seq tracks from triplicate MDA-MB-231 cells taken after either *RBMX* depletion (“RBMX siRNA”) or control treatment (“Control siRNA”). Previously identified polyadenylation (polyA) sites that are annotated in PolyASite 2.0 (54) are shown below the RNA-seq tracks. A putative novel termination sites within *BRCA2* that is preferentially used in RBMX-depleted cells is indicated with red arrows. **(B)** Pie chart representing the proportion of premature transcription termination events that occur upon RBMX depletion at sites already annotated in PolyASite 2.0 (54). **(C)** Snapshot from IGV browser (37) over *BRCA2* exons 10-12 showing merged RNA-seq tracks from MDA-MB-231 cells taken after either *RBMX* depletion (“RBMX siRNA”) or control treatment (“Control siRNA”), as well as a comparative track produced as ratio between RBMX siRNA and Control siRNA tracks using the bamcompare tool from deepTools3.5.0 (58). Specifically, positive (RBMX siRNA)/(Control siRNA) values are coloured in red while negative (RBMX siRNA)/(Control siRNA) values are coloured in blue. The putative polyadenylation signal identified within *BRCA2* exon 11 is highlighted in orange. **(D)** Upper panel, schematic representation of PCR products amplified from regions located either upstream or across the putative polyA site within *BRCA2* exon 11. Bottom panel, Representative capillary gel electrophoretograms showing RT-PCR analysis of the two PCR products from *BRCA2* exon 11. **(E)** Bar chart associated with (D) shows the relative expression of the full-length *BRCA2* isoform (encompassing the putative polyA site) compared to control PCR amplification (upstream the putative polyA site), using data from three biological replicates. The error bars show standard error. *, p-value < 0.5 as calculated by t-test by 2-tailed t-test assuming equal variance. **(F)** Upper panel, schematic representation of the region that displays drop in RNA-seq reads upon RBMX knock-down (termination window, TW). The region “before” TW is defined from transcription start site (TSS) to the 5’ edge of TW and the region “after” TW is defined from the 3’ edge of TW to the last polyadenylation signal (PAS) for all genes in Figure 5 – Source Data 1. Lower panel, dot plot representing the fold-change in RNA-seq read density between cells treated with RBMX siRNA and control siRNA over the regions before (red dots) and after (blue dots) TW. Black lines connecting the two groups of dots indicate the individual change between the two regions for all genes in analysis. A fold-change equal to 1 (RNA density in RBMX-depleted cells = RNA density in control cells) is indicated with dashed grey line as reference. ****, p-value < 0.0001 as calculated by Wilcoxon matched-pairs signed rank test. **(G)** formula for calculating Termination Index (TI) from RNA-seq reads before and after TW. **(H)** Boxplot analysis of TI for transcripts in Figure 5 – Source Data 1, averaged across three biological replicates for RNA-seq read densities measured from MDA-MB-231 cells treated with either control siRNA or siRNA against RBMX. Plot was produced using ggplot2 v.3.3.2 on R v.4.0.2 (41). Median TI for control cells is indicated with dashed grey line as reference.****, p-value < 0.0001 as calculated by Wilcoxon matched-pairs signed rank test. **(I)** Boxplot analysis as in (H) for the indicated samples analysed from HEK293 (26). Median TI for the first sample of control cells is indicated with dashed grey line as reference.****, p-value < 0.0001 and ns, non-significant as calculated by Wilcoxon matched-pairs signed rank test.

In order to better understand the impact of RBMX knock-down on premature transcription termination, we used the IGV genome browser (37), annotation on previously mapped polyA sites (54) and the bamcompare comparative track (58). We visually defined a termination window (TW) within these genes where RNA-seq tracks drop in RBMX knock-down compared to control. We then quantified RNA-seq reads upstream (“before”) and downstream (“after”) of the TW (Figure 5F, top panel). MDA-MB-231 cells depleted for RBMX revealed a dramatic and significant reduction of RNA density after the TW. This indicates defective production of full-length RNAs for this panel of genes after RBMX depletion (Figure 5F, bottom panel). To quantitate the contribution of RBMX on transcription termination we defined a Termination Index (TI) as the ratio between the RNA-seq density over the region downstream of TW and the region upstream of TW (Figure 5G). Smaller TI values are associated with lower RNA-seq density at the end of the gene, indicative of premature transcription termination. In agreement with our hypothesis, a Wilcoxon paired test showed that average TI over the genes identified by our analyses was significantly decreased in RBMX-depleted cells compared to control (Figure 5H, and Figure 5 – Source Data 1). Almost all 64 genes in this panel displayed significant reduction in TI across all biological replicates as measured by t-test with multiple test correction (Figure 5 – Figure Supplement 1A, Figure 5 – Source Data 1). This same analysis applied to published RNA-seq data (26) confirmed this novel RBMX role in suppressing upstream poly(A) sites in HEK293 cells, where depletion of RBMX resulted in a similar, significant reduction of RNA-seq reads after TW and reduced TI values compared to control (Figure 5I, Figure 5 – Figure Supplements 1B, C and Figure 5 – Source Data 1). However, the same was not observed after depletion of either of the m6A methyltransferases METTL3 and METTL14 (26) (Figure 5I, Figure 5 – Figure Supplements 1 B, C and Figure 5 – Source Data 1), showing that RBMX function in transcriptional termination is m6A-independent.

The above results reveal that RBMX contributes to full-length mRNA production by preventing early transcription termination within a subset of genes. Some of these premature termination events involve use of alternative upstream terminal exons, such as within the *ASPH* Aspartate Beta-Hydroxylase oncogene (59) (Figure 5 – Figure Supplement 2A), or upstream cryptic terminal exons, for example within the *ABLIM3* gene (Figure 5 – Figure Supplement 2B). However, a number of other premature termination events occurred at polyA sites localised within ultra-long exons in genes involved in replication fork activity. These ultra-long exons were found within *BRCA2* (4932nt long exon 11), *FANCM* (1905nt long exon 14) (Figure 5 – Figure Supplement 3A); and *GEN1* (RBMX represses an alternative upstream polyA sites within the terminal exon of the gene) (Figure 5 – Figure Supplement 3B). RBMX also enables full-length inclusion of ultra-long exons within genes involved in other aspects of genome stability. These include *RESF1* (Retroelement Silencing Factor 1, 5109nt long exon 4) that negatively regulates endogenous retroviruses (60) (61) (Figure 5 – Figure Supplement 3C); *ASPM* (*Abnormal spindle-like microcephaly-associated*) that is essential for normal mitotic spindle function (62) (63) (Figure 5 – Figure Supplement 3D); and *KNL1* (*kinetochore scaffold 1*) that is essential for spindle-assembly checkpoint signalling and for correct chromosome alignment (64) (Figure 5 – Figure Supplement 3E).

**Figure 5 – Figure Supplement 1.**
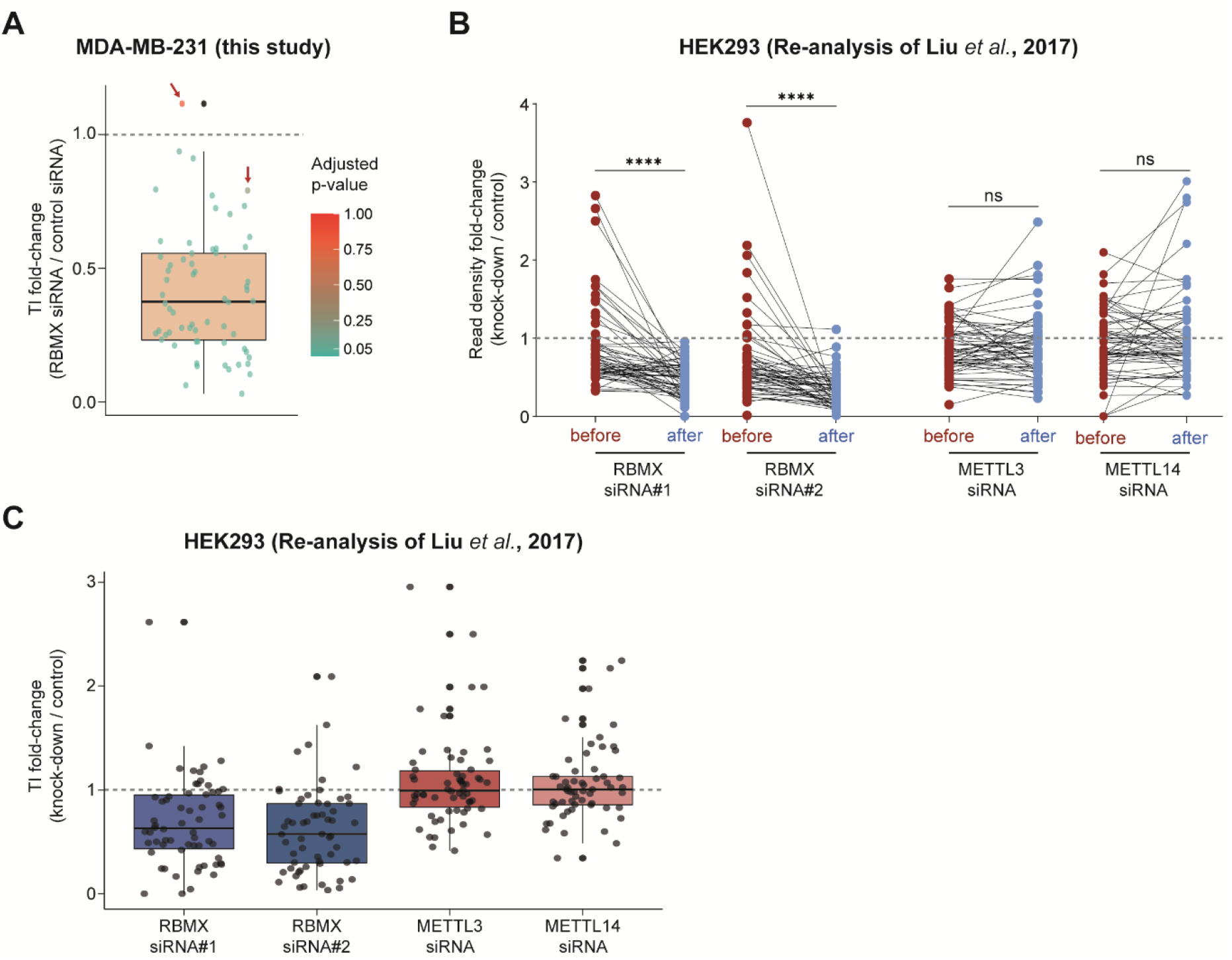
**(A)** Boxplot analysis shows the TI fold-change between cells treated with RBMX siRNA (TI_(RBMX siRNA)_) and cells treated with control siRNA (TI_(control siRNA)_). for all transcripts in Figure 5 – Source Data 1. TI_(RBMX siRNA)_/ TI_(control siRNA)_ ratios were calculated after averaging TI values from biological replicates for both samples. The overlapping jitter plot shows individual TI_(RBMX siRNA)_/ TI_(control siRNA)_ fold-change ratios. These are coloured according to the adjusted p-value (FDR method) calculated after t-test with multiple test correction across biological triplicates. The only two genes that show adjusted p-values above 0.05 are indicated with red arrows. Plot was produced using ggplot2 v.3.3.2 on R v.4.0.2 (41). **(B)** Dot plot analysis as in Figure 5F for the indicated samples analysed from HEK293 (26). ****, p-value < 0.0001 and ns, non-significant, as calculated by Wilcoxon matched-pairs signed rank test. **(C)** Boxplot analysis for TI fold-change was performed as in (A) for RNA-seq data from HEK293 (26). Jitter plot is shown as monochrome because t-test with multiple test correction could not be performed across the two only biological replicates provided from this dataset (26).

**Figure 5 – Figure Supplement 2.**
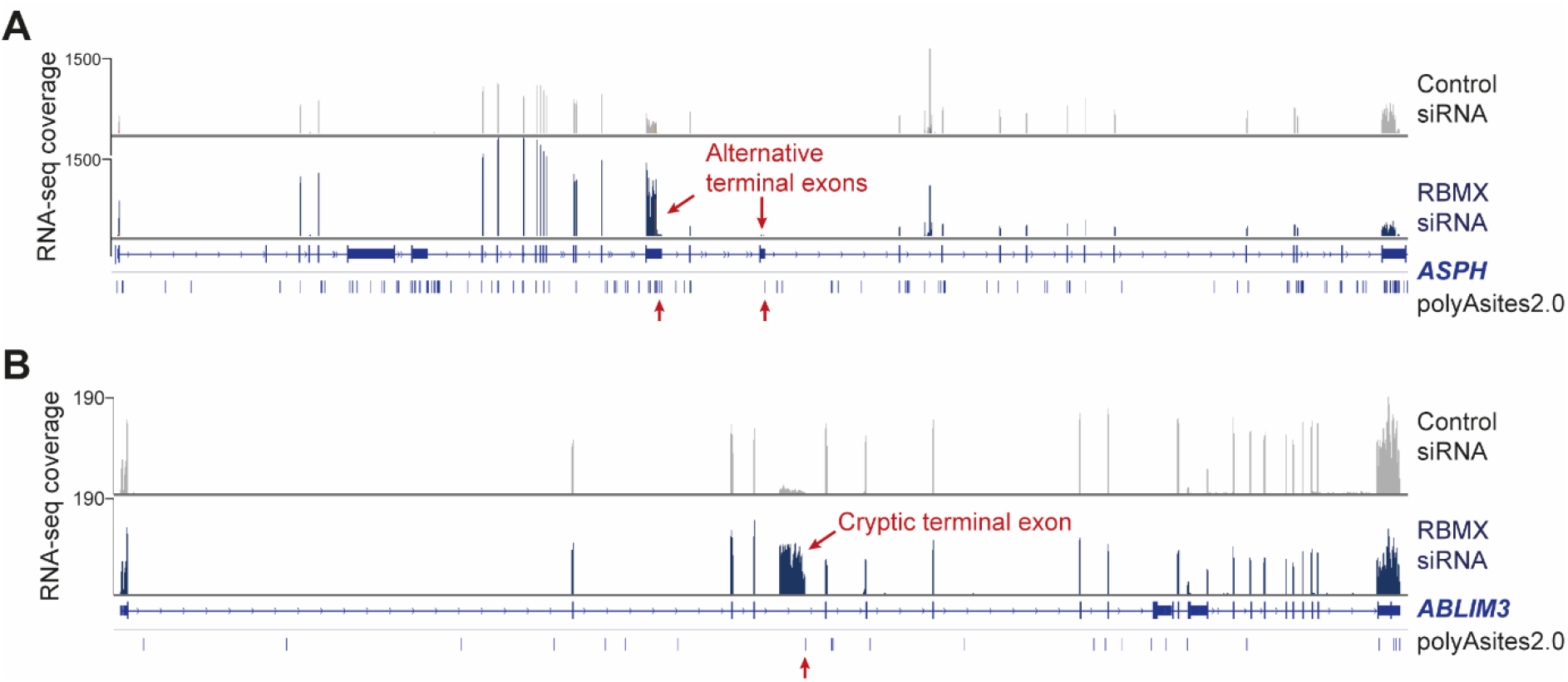
**(A-B)** Snapshots from IGV browser (37) over the *ASPH* (A) and *ABLIM3* (B) genes showing merged RNA-seq tracks from triplicate MDA-MB-231 cells taken after either *RBMX* depletion (“RBMX siRNA”) or control treatment (“Control siRNA”). Previously identified polyadenylation (polyA) sites annotated in PolyASite 2.0 (54) are shown for all tracks. Alternative upstream polyA sites within *ASPH* (A) and a cryptic upstream terminal exon within *ABLIM3* (D) preferentially used in RBMX-depleted cells are indicated with red arrows.

**Figure 5 – Figure Supplement 3.**
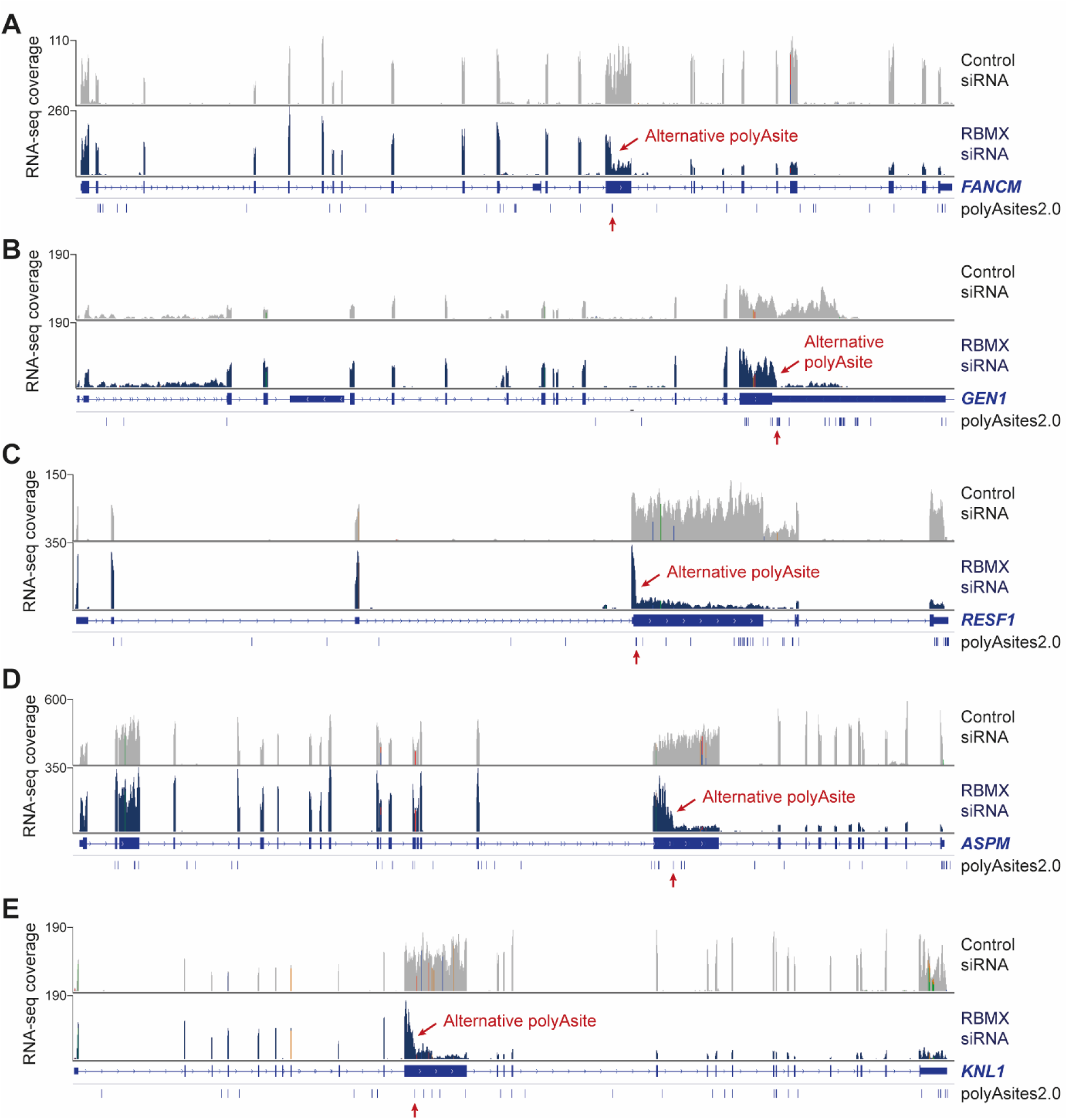
**(A-D)** Snapshots from IGV browser (37) over the *FANCM* (A), *GEN1* (B), *RESF1* (C), *ASPM* (D) and *KNL1* (E) genes showing merged RNA-seq tracks from triplicate MDA-MB-231 cells taken after either *RBMX* depletion (“RBMX siRNA”) or control treatment (“Control siRNA”). Previously identified polyadenylation (polyA) sites annotated in PolyASite 2.0 (54) are shown for all tracks. Alternative upstream polyA sites that are preferentially used in RBMX-depleted cells are indicated with red arrows.

### RBMX expression levels control gene networks involved in replication fork activity and DNA damage

The above data showed that RBMX is required for productive RNA processing of genes important for replication fork activity, including *ETAA1, REV3L, ATRX*, FANCM and *BRCA2*. However, depletion of RBMX in U2OS cells caused no defects in S phase of the cell cycle (21), and we observed a similar situation in MDA-MB-231 cells using Fluorescence-activated Cell Sorting (FACS) (Figure 6 – Figure Supplements 1A, B). Altogether, these results indicate that RBMX may modulate expression of other genes that can enable cell cycle progression to continue when the levels of key replication fork proteins drop. To further analyse this phenomenon, we examined the impact of RBMX depletion on the cellular transcriptome by analysing changes in RNA levels detected by our RNA-seq in MDA-MB-231 cells. Overall, 1596 genes showed an increased fold-change of at least 1.5 (log2FoldChange = 0.6), and 1691 showed a decreased fold-change of 0.65 (log2FoldChange = −0.6) or less of RNA levels in cells depleted for RBMX compared to control (adjusted p-value ≤ 0.05) (Figure 6A).

**Figure 6.**
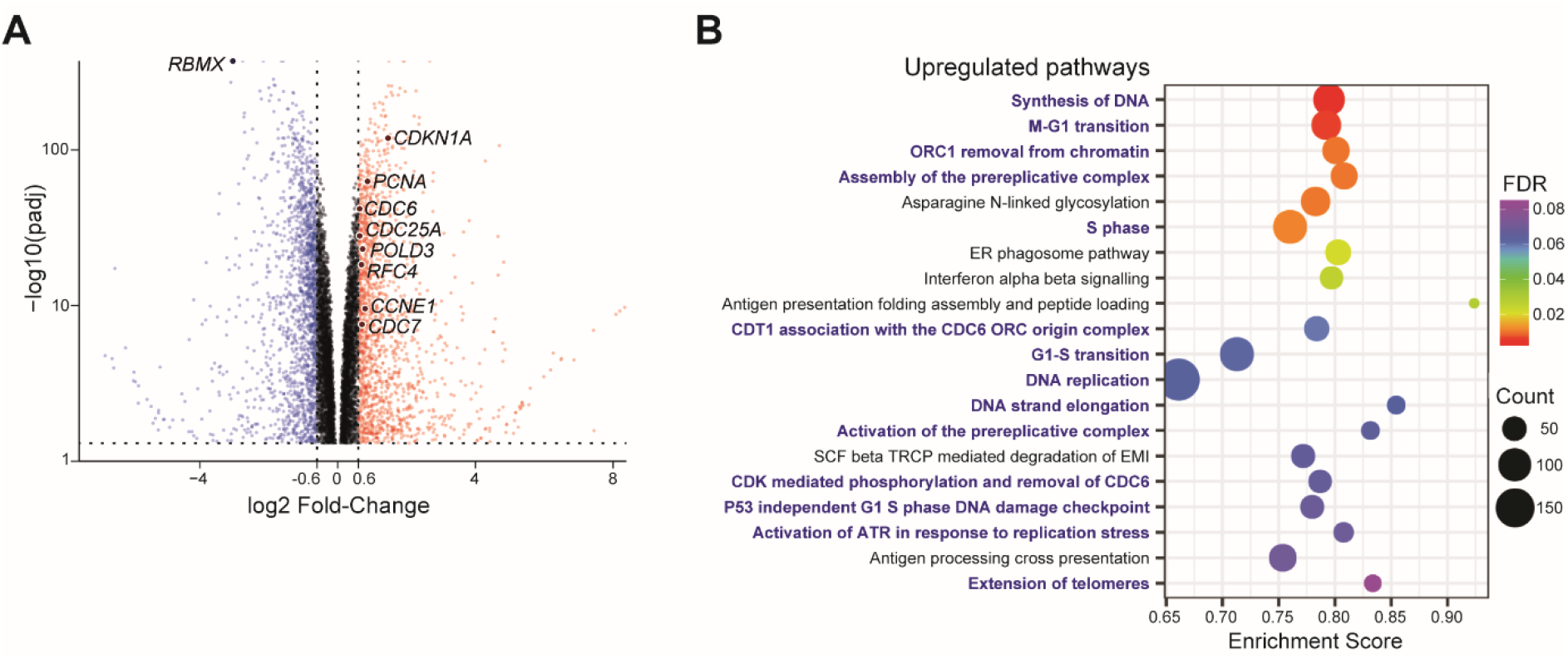
Depletion of RBMX from human breast cancer cells induces a gene expression signature related to DNA replication and repair. **(A)** Volcano plot showing changes in transcript levels after depletion of RBMX, as measured by RNA-seq. RNA-seq data were analysed using DESeq2 v.1.16.1 (43) and genes with either no RNA-seq reads or an adjusted p-value<0.05 were excluded. Blue dots, transcripts with a log2 Fold-Change between RBMX siRNA and control siRNA lower than −0.6 (Fold-Change < 0.66, *i.e*. at least 34% reduction). Red dots, transcripts with a log2 Fold-Change between RBMX siRNA and control siRNA higher than 0.6 (Fold-Change > 1.5, *i.e*. at least 50% increase). padj, adjusted p-value. Fold-changes in RNA levels over *RBMX* and genes discussed in the main text are highlighted. **(B)** Gene Set Enrichment Analysis (GSEA) was performed using the Broad Institute GSEA software (65) to identify the top 20 upregulated REACTOME pathways after RBMX depletion. Pathways involved in DNA replication are highlighted in blue. FDR, False discovery rate. Count, number of genes.

To identify patterns of gene expression that change on RBMX depletion we then used Gene Set Enrichment Analysis (GSEA), which takes into account fold-changes in RNA levels measured by RNA-seq (65). Strikingly, 14/20 of the most significant up-regulated pathways identified after RBMX depletion were involved in DNA replication and DNA damage response during S phase of the cell cycle, including activation of ATR in response to replication stress (Figure 6B) (15). Transcripts regulated by the E2F transcription factor have been reported to be maintained at high levels in response to replication stress (66). The gene sets up-regulated in response to RBMX depletion include some of these transcripts, specifically important regulators of S phase such as the cell division cycle proteins *CDC6, CDC7, CDC25A, CDC45*, the cyclins E1, D1 and D3, *CDKN1A* and *CDK4* (Figure 6 – Source Data 1). Moreover, other up-regulated genes encode important components of the replication machinery. These include the catalytic subunit of DNA polymerase epsilon (*POLE*) which is involved in chromosome replication and DNA damage repair; Replication Factor C2 and Replication Factor C4 (*RFC2* and *RFC4*) that are required for DNA elongation by DNA polymerases δ and ε; and the *POLD2, POLD3* and *PCNA* genes that encode proteins which increase activity of DNA polymerase δ, and help its recruitment to sites of DNA damage (Figure 6A and Figure 6 – Source Data 1). Importantly, none of these transcripts showed apparent mRNA processing defects upon depletion of RBMX. Overall, these data suggest that breast cancer cells can maintain cell cycle progression by subtly modulating gene expression patterns after RBMX depletion that relate to S phase of the cell cycle.

**Figure 6 – Figure Supplement 1.**
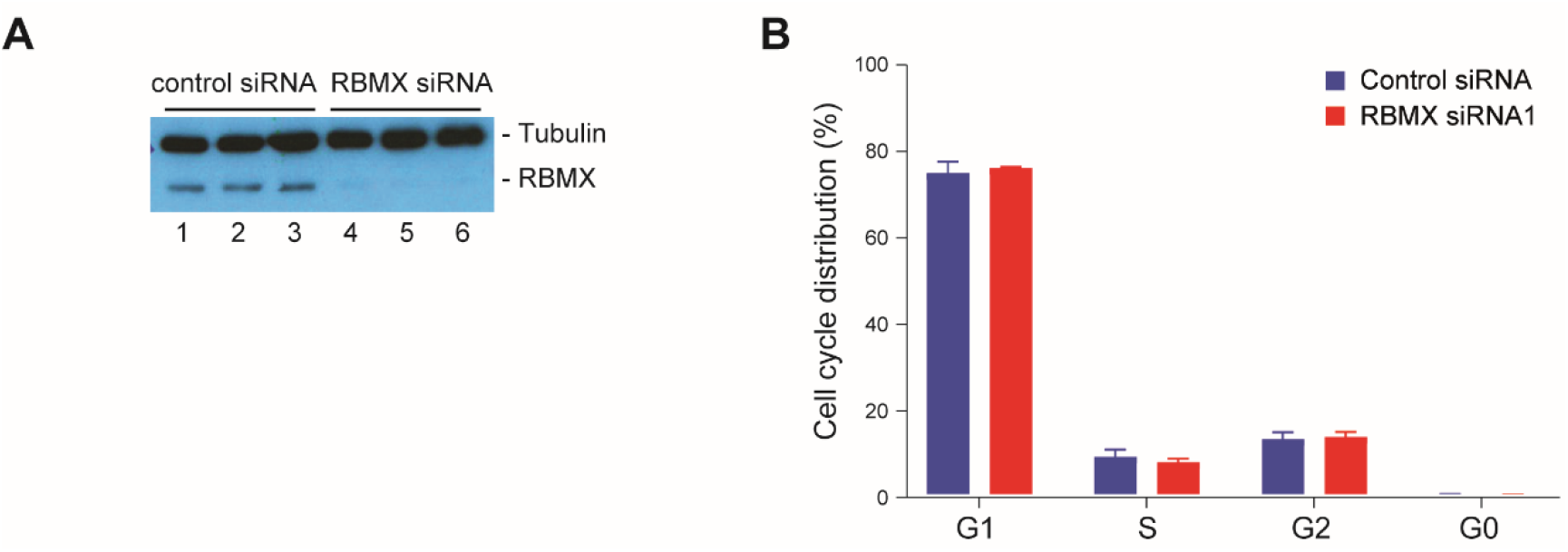
**(A)** Western blot analysis confirming reduction in RBMX protein levels in MDA-MB-231 cells after siRNA-mediated depletion of *RBMX*. Levels of RBMX protein were depleted around 90% when quantitated relative to tubulin. Cells from the same samples were analysed by flow cytometry in (B). **(B)** Flow cytometry analysis shows that siRNA-mediated depletion of *RBMX* causes no change in cell cycle distribution. Bars represent standard error from 3 biological replicates.

## Discussion

Here we have tested the hypothesis that RBMX controls genome stability via RNA processing. Supporting this, global analyses of RBMX-controlled mRNA processing patterns in human breast cancer cells show RBMX suppresses the use of splicing and polyadenylation sites within key genes that are crucial for genome stability (Figures 7A, B). This conclusion changes the way that we think about RBMX and DNA damage control, from a purely structural role at sites of replication fork stalling or DNA damage (15,21), to include an earlier role in gene expression patterns that regulate genome maintenance. Moreover, this better understanding of RBMX-controlled RNA processing patterns provides new molecular insights through which RBMX could operate as a tumour suppressor (4–8), and within gene expression networks in cancer cells (3).

**Figure 7.**
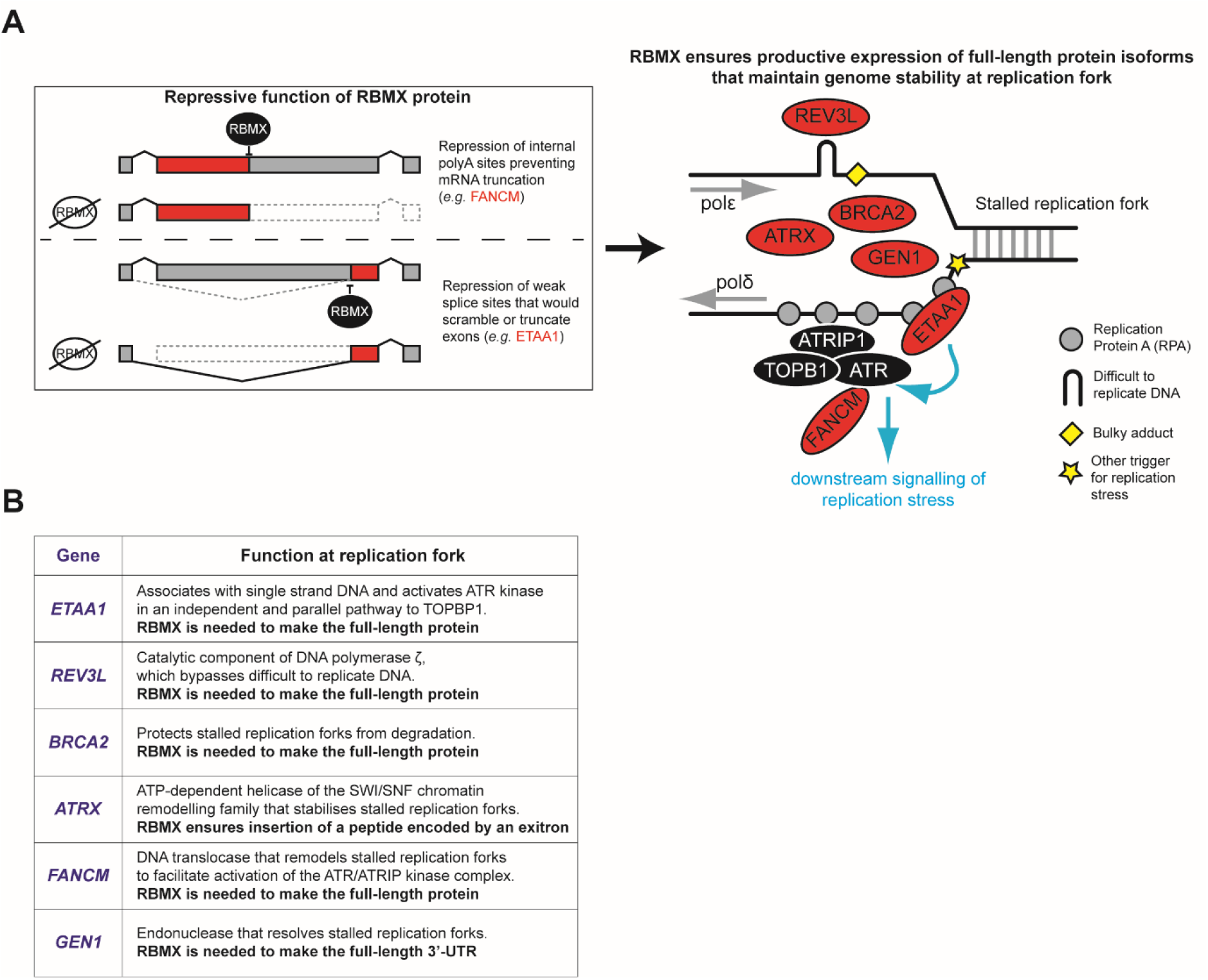
RNA processing by RBMX controls key genes for replication fork stability. **(A)** Schematic representation of the repressive function of RBMX in preventing use of cryptic mRNA processing sites thus promoting correct replication stress response and genome maintenance. **(B)** Schematic table indicating the function of RBMX-controlled genes shown in (A) at the replication fork.

The RBMX-regulated RNA processing events identified in this study have largely distinct properties compared with previous reported targets (26,27) in that they: (1) include a wider spectrum of RBMX-regulated events than just skipped exons (2); are largely suppressed by RBMX; and (3) seem to be regulated by RBMX largely independent of m6A RNA modification. The RNA processing defects detected in this study are conceptually similar to those detected in the mouse testis after the genetic deletion of the *RBMX* paralog *Rbmxl2* (34), which showed increased use of weak splice sites that would poison gene expression. Hence, although the actual regulated genes are different between human breast cancer cells and mouse testis, RBMX and Rbmxl2 share similar predominantly repressive activities that are important for productive gene expression.

Replication fidelity makes a key contribution to genome stability, and depletion of RBMX causes defective ATR activation in response to replication fork stalling (15). Our data here reveal that amongst the strongest defects in RNA processing patterns in response to RBMX depletion are six genes that encode key replication fork proteins (these are ETAA1, REV3L, BRCA2, ATRX, GEN1 and FANCM, Figures 7A, B). Most importantly, these include ETAA1 protein, which associates with single strand DNA at stalled replication forks, to activate ATR kinase in an independent and parallel pathway to TOPBP1 (17,45,46). ETAA1 is crucial for replication fork activity: cells directly depleted for ETAA1 protein (which also becomes virtually undetectable after RBMX protein depletion) become hypersensitive to replication stress and exhibit genome instability (45). RBMX is also required for productive expression of *REV3L*, which encodes the catalytic component of DNA polymerase ζ. This polymerase is used to by-pass sites of DNA adduct incorporation, or difficult to replicate DNA (51). The large exon in *REV3L* that is disrupted by RBMX depletion encodes a 1386 amino acid disordered peptide stretch important for efficient polymerase ζ activity, and inactivation of REV3L causes genomic instability (51). Furthermore, RBMX represses an exitron within *ATRX*, a gene encoding a protein that stabilises stalled replication forks (67). RBMX is similarly important for full-length expression of the *FANCM* gene, which encodes a DNA translocase that remodels stalled replication forks to facilitate activation of the ATR/ATRIP kinase complex (68). RBMX promotes full-length UTR expression from the *GEN1* gene, which encodes a protein that resolves stalled replication forks (69). RBMX is also critical for full-length expression of the *BRCA2* gene that protects stalled replication forks from degradation (70,71). Finally, we detect a subtle upregulation of other genes involved in DNA replication and DNA damage control after RBMX depletion, which likely represents a cellular response to increased DNA replication fork stalling (66). Previous studies have shown that Chk1 kinase inhibits the E2F6 transcriptional repressor to promote upregulation of cell-cycle transcriptional programmes in response to replication stress (66). However, further analysis will be required to clarify the mechanisms of compensation that maintain replication fork stability in the absence of RBMX. We also cannot exclude that shorter proteins are made from truncated mRNAs after RBMX-depletion that might interfere with the function of the full-length protein isoforms.

Although RBMX has previously been associated with splicing regulation, the biggest group of RNA processing events that we identified as repressed by RBMX are upstream polyA sites and alternative upstream terminal exons. The selection of polyA sites and alternative splice sites both depend on assembly of RNA protein complexes, and are driven by recognition of consensus sequences within pre-mRNAs. RBMX is part of the hnRNP family of proteins, many of which coat RNA and so may block RNA processing signals within pre-mRNAs (72). RBMX binding to pre-mRNA may hence act as a general signal to block use of aberrant RNA processing sites embedded in transcripts. Current data are consistent with a model by which RBMX could facilitate splicing inclusion and full-length transcription of important protein coding genes by sterically masking cryptic mRNA processing sites (Figure 7A). This model is supported by the presence of multiple RBMX binding sites (26) across several of these large exons included within *REV3L, ATRX, RIF1, ASPM*. RBMX may also operate in conjunction with other RNA binding proteins to bind RNA. This latter mechanism might be important for *ETAA1* exon 5. Although Tra2β associates to *ETAA1* RNA near the cryptic splice site within exon 5 (see Supplementary Figure 4), analysis of published PAR-CLIP (26) revealed that RBMX might not do the same. Co-transfection experiments indicate that the RBMX relies on Tra2β binding to RNA to correctly process *ETAA1* mRNA. One intriguing possibility might be that RBMX interacts with Tra2β bound to RNA, creating a complex that sterically prevents the spliceosome to access the internal 3’ splice site within *ETAA1* exon 5.

Our data also provide insight into how ultra-long exons are processed. In mammals, exons are recognised by a process called exon definition, in which U1 and U2 snRNPs of the spliceosome interact with the 5’ and 3’ splice sites, and are stabilised by interactions with nuclear RNA binding proteins attached within and nearby to exon sequences (73). The median size of human exons is approximately 129 base pairs (bp), however a large number of genes contain exons that can reach several kilobases (kb) in length. To what extent specific mechanisms exist to allow the spliceosome to recognize splice sites that are far apart, thus enabling splicing inclusion of ultra-long exons, is not well understood. Our data here indicate that RBMX plays a key role in suppressing the use of aberrant RNA processing signals that would interrupt the inclusion of long exons. Long exons might be particularly destabilised by loss of RBMX, since their increased length would make it more likely to contain short sequence motifs that could be utilised by other RNA processing pathways. Moreover, deficiencies in proper RNA processing pathways of long exons might in contribute to developmental defects and human disease associated with RBMX deficiency. In fact, *RBMX* is mutated in the human X-linked mental retardation Shashi syndrome (14), and individual mutation of genes that rely on RBMX for productive expression can cause mental retardation (*ATRX*) (74) microcephaly (*KNL1* and *ASPM*) (75,76). Consistent with this hypothesis, loss of RBMX also affects brain development in zebrafish (13).

## Materials and methods

### Cell culture and cell lines

MDA-MB-231 (ATCC^®^ HTB-26™), MCF7 (ATCC^®^ HTB-22™), and HEK293 (ATCC^®^ CRL-1573) cells were grown as previously described (48). Cell line validation was carried out using STR profiling according to the ATCC^^®^^ guidelines. All cell lines underwent regular mycoplasma testing.

### siRNA knockdown

RBMX transient knockdown was established using two different siRNAs targeting *RBMX* mRNA transcripts (hs.Ri.RBMX.13.1 and hs.Ri.RBMX.13.2, from Integrated DNA Technologies). Negative control cells were transfected with scramble siRNA (IDT). Cells were seeded onto 6-well plates at a confluence of approximately 1×10^6^ and incubated for 24hours. After the incubation period, 6μl of 10μM siRNA was diluted in 150μl of Opti-MEM which was then mixed with 6μl of Lipofectamine RNAiMAX also diluted in 150μl. The combined reaction mix was incubated at room for 5 min and then added dropwise onto the seeded cells. Transfected cells were then incubated for 72h at 37°C before harvesting.

### RNA-seq

RNA was extracted from cells using RNeasy Plus Mini Kit (Qiagen) following manufacturer’s instructions and re-suspended in nuclease-free water. RNA samples were DNase treated (Invitrogen). Pair-end sequencing was done initially for two biological samples, one of negative control siRNA treated MDA-MB-231 cells and one of RBMX siRNA treated cells, using an Illumina NextSeq 500 instrument. This RNAseq data is deposited at GEO (accession GSE158770). Adapters were trimmed using trimmomatic v0.32. Three additional biological repeats of negative control and RBMX siRNA treated MDA-MB-231 cells were then sequenced using an Illumina HiSeq 2000 instrument. The base quality of raw sequencing reads was checked with FastQC (77). RNA-seq reads were mapped to the human genome assembly GRCh38/hg38 using STAR v.2.4.2 (78) and subsequently quantified with Salmon v. 0.9.1 (79) and DESeq2 v.1.16.1 on R v.3.5.1 (43). All snapshots indicate merged tracks produced using samtools (80) and visualised with IGV (37) unless specified.

### Identification of splicing changes

Initial comparison of single individual RNA-seq tracks from RBMX-depleted and control cells was carried out using MAJIQ (36), which identified 596 unique local splicing variations (LSV) at a 20% dPSI minimum cut off from 505 different genes potentially regulated by RBMX.

These LSVs were then manually inspected using the RNA-seq data from the second RNA sequencing of biological replicates for both RBMX-depleted and control cells, by visual analysis on the UCSC browser (81) to identify consistent splicing changes that depend on RBMX expression. The triplicate RNA-seq samples were further analysed for splicing variations using SUPPA2 (35), which identified 6702 differential splicing isoforms with p-value < 0.05. Predicted splicing changes were confirmed by visual inspection of RNA-seq reads using the UCSC (81) and IGV (37) genome browsers. Splice site strength at *ETAA1* exon 5 were calculated using MaxEntScan::score3ss (http://hollywood.mit.edu/burgelab/maxent/Xmaxentscan_scoreseq_acc.html)

### RNA extraction and cDNA synthesis for transcript isoform analysis

RNA was extracted using standard Trizol extraction protocol and DNAse treated using DNA-free kit (Invitrogen). The RNA from siRNA-treated cells was extracted using standard Trizol RNA extraction (Life Technologies) following manufacturer’s instructions. cDNA was synthesized from 500 ng total RNA in 10 μl reactions using Superscript VILO cDNA synthesis kit (Invitrogen) following manufacturer’s instructions. To analyse the splicing profiles of the alternative events primers were designed using Primer 3 Plus and the predicted PCR products were confirmed using the UCSC *In-Silico* PCR tool. *ETAA1* transcript isoform containing the long exon 5 was amplified by RT-PCR using primers 5’-GCTGGACATGTGGATTGGTG-3’ and 5’-GTGCTCCAAAAAGCCTCTGG-3’, while *ETAA1* transcript isoform containing the short exon 5 was amplified using primers 5’-GCTGGACATGTGGATTGGTG-3’ and 5’-GTGGGAGCTGCATTTACAGATG-3’. RT-PCR with this second primer pair could in principle amplify also a 2313 bp product from the *ETAA1* transcript isoform containing the long exon 5, however PCR conditions were chosen to selectively analyse shorter fragments. *BRCA2* transcript isoform encompassing the putative polyA site within exon 11 and a control fragment upstream this site were amplified by multiplex RT-PCR using a forward primer 5’-TCAGGTAGACAGCAGCAAGC-3’ and two reverse primers, respectively 5’-TCCCTCCTTCATAAACTGGCC-3’ and 5’-AACCCCACTTCATTTTCATCTGTT-3’. All PCR reactions were performed using GoTaq^®^ G2 DNA polymerase kit from Promega. All PCR products were examined using the QIAxcel^®^ capillary electrophoresis system 100 (Qiagen).

### Western blot analyses

Harvested cells treated with either control siRNA or siRNA against RBMX were resuspended in 100mM Tric HCL, 200mM DTT, 4% SDS, 20% Glycerol, 0.2% Bromophenol blue, then sonicated and heated to 95°C for 5 minutes. Protein separation was performed by SDS-page on a 10% acrylamide gel. Proteins were then transferred to a nitrocellulose membrane, incubated in blocking buffer (5% Milk in 2.5% TBS-T) and stained with primary antibodies diluted in blocking buffer to the concentrations indicated below, at 4°C over-night. After incubation the membranes were washed three times with TBS-T and incubated with the secondary antibodies for 1 hour at room temperature. Detection was carried out using the Clarity™ Western ECL Substrate (GE Healthcare Systems) and developed using medical X-ray film blue film in an X-ray film processor developer. The following primary antibodies were used: anti-RBMX (Cell Signalling, D7C2V) diluted 1:1000, anti-SGO2 (Bethyl Laboratories, A301-261A) diluted 1:1000, anti-ETAA1 (Sigma, HPA035048) diluted 1:1000, anti-Tubulin (Abcam, ab18251) diluted 1:2000 and anti-beta-actin (Abcam, ab5441) diluted 1:1000.

### Minigene Experiments

A genomic region containing the weak 3’ splice site within *ETAA1* exon 5 and flanking sequences were PCR amplified from human genomic DNA using the primers ETAA11295F (5’-AAAAAAAAACAATTGGAACATGGAGCCAAACTAACTC-3’) and ETAA11295R (5’-AAAAAAAAACAATTGTGATAGAATGGAGACTTGGGGA-3’), and cloned into pXJ41 (47). Splicing patterns were monitored after transfection into HEK293 cells with expression constructs encoding GFP, RBMX-GFP, RBMXL2-GFP, or Tra2β-GFP or deletion variants of the above plasmids as previously described (32,50). Splicing analysis was carried out in HEK293 cells after lipofectamine 2000 (Invitrogen) transfection of plasmids. RNA was extracted with TRIzol (Invitrogen), and analysed using a one-step RT-PCR (PCR with reverse transcription) kit from Qiagen, both using the standard protocol. RT-PCR experiments used 100 ng of RNA in a 5-μl reaction using a multiplex RT-PCR using primers: 5’-GCTGGACATGTGGATTGGTG-3’, 5’-GTGGGAGCTGCATTTACAGATG-3’ and 5’-GTGCTCCAAAAAGCCTCTGG-3 ‘. Reactions were analysed and quantified by capillary gel electrophoresis.

### Transcription termination analyses

Termination widows (TW) for all genes in Figure 5 – Source Data 1, which appear prematurely terminated upon treatment with RBMX siRNA, were defined as the regions where RNA-seq reads drop on tracks from RBMX-depleted cells but not on tracks from control cells. Confirmation of TW was carried out by visual inspection of a comparative alignment track generated using the bamcompare tool from deepTools v3.5.0 (58) on IGV browser (37). Subsequently, a .SAF annotation file was built to define the regions *“before”* and *“after”* TW for each premature transcription termination event. Specifically, regions *“before”* were defined from Transcription Start Site (TSS) to TW Start coordinate and regions *“after”* were defined from TW End coordinate to Gene End coordinate. Strand (+/-), TSS and Gene End annotations were obtained from UCSC (81). The .SAF file was used as index to quantify RNA-seq reads from .BAM files using the featureCounts tool from Subread package v.2.0.1 (82). Quantification was performed on biological triplicates for RNA-seq tracks from MDA-MB-231 (this study) and on merged .BAM files from biological duplicates for RNA-seq tracks from HEK293 (26). RNA-seq densities *“before”* TW and *“after”* TW were averaged, and the statistical difference between RNA-seq densities over the two regions was calculated by Wilcoxon matched-pairs signed rank test (paired, non-parametric) after testing for normal distribution using GraphPad Prism v.8.2.1 for Windows (GraphPad Software, San Diego, California USA, www.graphpad.com) for all samples from MDA-MB-231 (this study) and HEK293 (26). Termination Index (TI) was calculated as *(RNA-seq density “after”)/(RNA-seq density ”before”)* for all events. Statistical significance of TI changes in RBMX knock-down compared to control was calculated by t-tests with multiple test correction using the qvalue package on R v.4.0.2 (83) across biological triplicates from MDA-MB-231 cells (this study), validating 62/64 transcription termination events (Figure 5 – Source Data 1). The same test could not be performed on TI calculated from HEK293 RNA-seq, as only two biological replicates are present in this dataset (26). For this reason, TI values were averaged across biological replicates for RBMX siRNA and control siRNA treated MDA-MB-231 cells, and statistical significance of TI changes upon RBMX depletion was calculated again by Wilcoxon matched-pairs signed rank test (paired, non-parametric) after testing for normal distribution using GraphPad Prism v.8.2.1 for Windows (GraphPad Software, San Diego, California USA, www.graphpad.com). The same test was performed on TI values calculated from merged RNA-seq tracks from HEK293 (26). Finally, TI fold-change ratios were calculated as *averageTI(RBMX siRNA)/averageTI(control siRNA)*. All TI fold-change ratios below 1 (63/64) confirmed reduction of RNA-seq reads after TW in RBMX knock-down compared to control (Figure 5 – Source Data 1). Similar TI fold-change ratios were calculated for HEK293 cells treated with either of two RBMX siRNAs, METTL3 siRNA and METTL14 siRNA from (26) over their respective control siRNA.

### Gene ontology analyses

Gene Ontology Enrichment Analyses shown in dot-plots and chord-diagram were performed with GOstats v.2.54.0 (40) with a p-value cut-off of 0.05, using the Bioconductor annotation data package org.Hs.eg.db v.3.11.4 and the whole list of genes detected by RNA-seq in MDA-MB-231 (i.e. excluding genes with no RNA-seq reads in the control samples) as background universe. P-values were adjusted by hypergeometric test first, and then by false discovery rate method on R v.4.0.2 (83). The chord diagram was produced for terms with count > 4 and size < 250 using the GOplot (v1.0.2) (42). The dot-plots were produced using ggplot2 v.3.3.2. (41) focussed on representative terms that had adjusted p-value (FDR) < 0.05.

Gene Set Enrichment Analysis (GSEA) was performed using the Broad Institute GSEA software v.3.0 (65). Genes identified by RNA-seq were ranked using log10(p-value) with a negative sign for down-regulated genes and positive sign for up-regulated genes. Enrichment was queried for REACTOME pathways using the pre-ranked tool of GSEA software with 1000 permutations.

### Analysis of long human exons

Annotations of all human exons related to position and size were downloaded from Ensembl Genes 101 (http://www.ensembl.org/biomart/). Density plot was created using ggplot2 v.3.3.2. (41) on R v.4.0.2.

### FACS

Three biological replicates of MDA-MB-231 cells treated with either control siRNA or siRNA against RBMX were washed in PBS and fixed in 70% ethanol. Cells were permeabilised using 0.1% Triton X-100 (Sigma) in PBS, then stained with 1:100 cy5-coupled MPM2 (Merck/Millipore), treated with 0.2mg/ml RNAseA (Thermo Scientific) and stained with 50 μg/ml propidium iodide (Invitrogen) for 20 minutes before analysis. Samples were analysed for DNA content by flow cytometry on a BD LSRFortessa™ cell analyser. Cell cycle distribution was calculated after appropriate gating of cell populations in FL-2-Area vs FL-2-Width plot of PI fluorescence.

## Acknowledgements

We thank Professor Nicola Curtin and Dr. Louise Reynard for comments on the text; and Luisa Thomas Eke, Scott Naylor, Hannah Power, Rhiannon Cook, Alexa Coopersmith and Chan Sin Min for their contributions to the project.

